# RiboWin: A computational framework to identify non-canonical open reading frames

**DOI:** 10.64898/2026.10.01.752299

**Authors:** D Vaibhav, N Aniruddha, Yadav K Bhavana, Sushant Phadnis, Bhupendra Verma, YS Ravikumar

## Abstract

**Background:** Ribosome profiling is widely used to map translation across the genome, yet current sORF discovery pipelines are constrained by their reliance on annotated protein coding regions and by treating biological replicates primarily as tools for quantification rather than as strict filters for reproducibility. This results in two major issues, the systematic exclusion of pseudogene derived loci from analysis and the generation of extremely large candidate sets in which true translation events are difficult to distinguish from noise. To overcome these limitations, a five-stage framework was developed that expands the search space to include pseudogene loci while enforcing dual biological replicate concordance as a hard binary filter.

**Results:** When applied to four human cell lines with matched replicates and benchmarked against existing tools, the framework produces dramatically smaller and more reliable candidate sets, reducing thousands to millions of predicted sORFs down to a small number of high confidence concordant loci. Across HeLa cell cycle conditions, a limited set of unique concordant sORFs was identified spanning multiple transcript classes including upstream ORFs, retained introns, and nonsense mediated decay transcripts, along with a pseudogene derived candidate. Notably, the pseudogene encoded sORF RPL26P19 was detected consistently in HeLa cells but missed or inconsistently identified by existing tools, while a separate pseudogene sORF from UBE3AP2 was observed specifically in normal primary cell types and absent in HeLa. Both candidates were later found to be supported by entries in OpenProt, even though this was not known during the analysis, which gives added confidence that the results are genuine and not biased by prior expectations.

**Conclusions:** Overall, these findings suggest that pseudogenes are not just inactive remnants in the genome but can act as real, functional sources of translated sORFs in a cell type specific manner. At the same time, it becomes clear that most current tools are not designed to detect these signals effectively. This highlights the need for approaches that both broaden the regions being searched and apply stricter reproducibility checks, so that truly meaningful translation events can be identified more reliably.

## Background

Ribosome profiling (Ribo-seq), introduced by [1] , has fundamentally transformed our view of the translated genome by providing nucleotide-resolution snapshots of ribosome occupancy across the transcriptome. By capturing ribosome-protected mRNA fragments (RPFs) of approximately 28 to 30 nucleotides, the technique offers a direct readout of translational activity at genome-wide scale. Its application to small open reading frame (sORF) discovery has been particularly transformative. sORFs encode peptides of up to 100 amino acids and reside in regions previously dismissed as non-coding ([2] ; [3] ; [4] ). Large-scale screens have catalogued tens of thousands of candidates translated regions from single datasets ([5] ; [6] ;[7] ; [8] ), and dedicated analysis tools including RiboCode ([9] , RibORF [10], and Ribo-TISH [11] have been developed to systematise this discovery process.

A fundamental challenge confronts the field; however, ribosome occupancy alone does not prove translation. Scanning ribosomes, stalled elongation complexes, and non-productive initiation events can generate Ribo-seq footprint signals that may be difficult to distinguish from productive translation in a single experiment ([12] ; [13] ; [5] ). The scale of this problem is not subtle. Applied to asynchronous HeLa cells in this study, Ribo-TISH identified 842,615 candidates in replicate 1 and 2,167,749 in replicate 2, a 2.57-fold discrepancy between two independent replicates from the same biological condition, with only 12.63% of replicate 1 candidates shared. A systematic multi-tool comparison using the same data found approximately 2% agreement across three major sORF tools, making clear that the overwhelming majority of candidates from any single tool are not reproducible. The most straightforward remedy is to require that candidates independently pass all filters in two biological replicates before being reported; yet this is conspicuously absent from the standard workflow. Biological replicate concordance is considered non-negotiable in essentially every other quantitative genomics context ([14] ; [15] ), yet in sORF discovery replicates are almost universally used for downstream quantification rather than as a binary quality gate at the candidate selection stage.

Pseudogenes represent a particularly compelling but neglected target for sORF discovery. The human genome harbours an estimated 14,000 to 20,000 pseudogenes ([16] ; [17] ), virtually all of which are absent from standard protein-coding annotation and therefore outside the search space of every existing sORF tool. This exclusion rests not on evidence of non-translation but on the historical assumption that pseudogenes are functionally inert which is an assumption that has been steadily eroding. Pseudogenes were historically considered non-functional, but this view has been challenged by studies demonstrating regulatory roles of pseudogene transcripts, including competitive endogenous RNA activity and transcriptional control ([18] ; [19]). In parallel, ribosome profiling studies have revealed widespread translation outside annotated coding regions, including within non-canonical transcripts, with some evidence suggesting translation may also occur in pseudogene-derived RNAs ([6] ; [5] ). Beyond the well-characterised AUG start codon, ribosome footprint mapping has revealed that eukaryotic translation initiates from non-AUG codons far more frequently than previously appreciated, generating functionally important protein products under specific cellular contexts [20] . The complexity of translation regulation extends beyond initiation codons. Ribosomal RNA-derived fragments (rRFs), a class of non-coding RNAs generated from rRNA, have emerged as active participants in translational control, regulating translation in a species-specific manner across diverse biological contexts [21] . Similarly, epitranscriptomic modifications such as m6A and pseudouridine directly influence mRNA stability, ribosome recruitment, and translational efficiency, adding a further layer of complexity to the determination of whether a given transcript is productively translated [22] . Ribosomal protein pseudogenes are of particular interest, their parent genes are among the most abundantly translated in the cell, and pseudogene copies may retain Kozak context, codon optimality, and translational regulatory elements from their progenitors. Whether they produce translated products detectable by a high-stringency Ribo-seq framework has not been rigorously tested.

Despite these observations, systematic identification of reproducibly translated pseudogene derived sORFs remains limited. A parallel gap exists within existing alternative protein databases, OpenProt catalogues AUG-initiated ORFs as AltProt entries, yet the majority carry no experimental support ([23]; [24] ), representing computational forecasts that rigorous Ribo-seq evidence can directly validate. The lack of experimental confirmation for many database entries underscores the need for pipelines that can provide high-confidence, reproducible evidence for novel translation events at non-canonical loci.

The cell cycle provides a biologically powerful and underutilised framework for assessing the specificity of sORF translation. Beyond the cell cycle, translation regulation is also central to host-pathogen interactions, where RNA-binding proteins orchestrate the translation of both viral and host mRNAs [25,26] . These studies demonstrate that translation is not a static process but is dynamically regulated in response to diverse cellular contexts, reinforcing the need for rigorous, context-aware detection of translation events. Translational activity is dynamically regulated across the cell cycle, with well-characterised suppression of cap-dependent translation during mitosis and phase-specific modulation of translation programs [27] ; [28]. A sORF that reproducibly translates in asynchronous or M phase conditions but is absent in S phase provides considerably stronger evidence of genuine biological regulation than one detected only in asynchronous populations. No previous study has systematically applied stringent replicate-concordance filtering to compare the reproducibly translated non-canonical proteome across asynchronous, S phase, and M phase conditions within a single cell type. This gap motivated the design of the current framework.

Furthermore, the field has long grappled with the challenge of distinguishing genuine microproteins from noise. Several independently validated microproteins have been discovered in transcripts previously classified as non-coding. [4] identified myoregulin as a micropeptide encoded by a transcript previously considered a long non-coding RNA, demonstrating its direct role in muscle performance by regulating the SERCA pump. Similarly, DWORF was discovered as a positive regulator of SERCA activity [29] ; [30] identified the HOXB-AS3 peptide, which suppresses colon cancer growth by directly inhibiting pyruvate kinase M2. These discoveries established a precedent that biologically functional microproteins can be encoded in unexpected genomic locations, strengthening the rationale for comprehensive, pseudogene-inclusive sORF screening frameworks.

The aims of this study were: (a) to develop a Ribo-seq sORF discovery framework with a pseudogene-inclusive search space and mandatory dual biological replicate concordance; (b) to detect actively translated sORFs by analyzing ribosome profiling (Ribo-seq) data and applying translation evidence filters such as ribosome footprint and frame periodicity; (c) and to evaluate the translation efficiency of sORFs across different cell cycle conditions of HeLa (Asynchronous, S phase, and M phase), fibroblast, keratinocytes, and melanocytes using RNA-seq integration. Collectively, these aims address fundamental gaps in the current state of sORF discovery which are, insufficient search space breadth, absence of mandatory reproducibility filters, and the neglect of pseudogene-derived translation as a biological phenomenon.

## Methods

### Datasets

All datasets used in this study are available in NCBI Gene Expression Omnibus (GEO). HeLa cell Ribo-seq and their corresponding RNA-seq datasets were obtained for three cell cycle conditions, each with two biological replicates. Cell cycle synchronisation in the used dataset was achieved using double thymidine block for S phase arrest and nocodazole treatment for M phase arrest, with asynchronous cells serving as the unsynchronised reference condition. RNA-seq accession numbers for HeLa are asynchronous replicates 1 and 2 (GSM2100595, GSM2100594), M phase replicates 1 and 2 (GSM2100591, GSM2100590), and S phase replicates 1 and 2 (GSM2100588, GSM2100587). Ribo-seq accession numbers for HeLa are asynchronous replicates 1 and 2 (GSM2100602, GSM2100601), M phase replicates 1 and 2 (GSM2100598, GSM2100599), and S phase replicates 1 and 2 (GSM2100596, GSM2100597).

For the multi cell-line analysis, publicly available Ribo-seq and matched RNA-seq datasets were obtained for three primary human cell lines, each with two biological replicates. RNA-seq accession numbers are fibroblast replicates 1 and 2 (GSM7430304, GSM7430303), keratinocyte replicates 1 and 2 (GSM7430302, GSM7430301), and melanocyte replicates 1 and 2 (GSM7430300, GSM7430299). Ribo-seq accession numbers are fibroblast replicates 1 and 2 (GSM7430296, GSM7430295), keratinocyte replicates 1 and 2 (GSM7430294, GSM7430293), and melanocyte replicates 1 and 2 (GSM7430292, GSM7430291). The choice of these three primary cell lines (fibroblast, keratinocyte, and melanocyte) enables a cancer versus normal comparison across cell types with distinct developmental origins and translational programs.

### RiboWin GitHub repository

The repository was organized into a modular and reproducible multi-phase workflow named RiboWin for the identification and validation of translated (sORFs) using RNA-seq and Ribo-seq data [31] . The framework architecture separates each analytical stage into dedicated directories and executable scripts to ensure scalability, reproducibility, and ease of maintenance. The workflow is supported by centralized configuration files (config.yaml and samples.tsv), while raw datasets and sequencing files are maintained within the data/ directory. Candidate sORFs are first identified using sorf_discovery.py, filtered and deduplicated through sorf_cleaning.py, and converted to BED format via make_bed.py. RNA-seq datasets are then processed using run_phase2.sh to generate transcript abundance and reference expression profiles, while Ribo-seq datasets undergo preprocessing, alignment, P-site calibration, and assignment through run_phase3.sh, psite_calibration.py, and psite_assignment.py. Ribosome occupancy data are subsequently integrated with coding sequence annotations via stage4_translation.py, and triplet periodicity analysis with replicate concordance filtering is applied through stage5_periodicity.py and merge_reps.py to retain high-confidence translated sORFs. Finally, translation efficiency is computed using TE_compute.py and filtered through TE_filter.py to identify TE-associated translated candidates.

It also includes workflow testing (run_test_quick.sh and run_test_full.sh) for validation. The complete pipeline is executed through the master orchestration script run_all.sh within a Conda-managed environment specified in environment.yml.

### Read Processing and Alignment

Quality control and trimming of raw FASTQ data were performed with Trimmomatic (version 0.39) with default settings. Sequences that were less than 25 nucleotides after trimming were removed from further analysis. Quality-controlled reads were mapped against the human reference genome (hg38/GRCh38) using STAR (version 2.7.10a) with the following arguments: --outSAMtype BAM SortedByCoordinate, --outFilterMismatchNmax 2, --alignIntronMax 1000000, and -- alignMatesGapMax 1000000. The Ensembl release 109 gene annotation (GRCh38) was used as the genome annotation for STAR alignment and transcript-level ORF enumeration. The choice of a maximum of two mismatches per alignment is consistent with standard Ribo-seq alignment practices and represents a balance between mapping sensitivity and specificity. Uniquely mapped reads were identified using the NH tag in the BAM alignment files, and reads with MAPQ scores below 10 were excluded from downstream P-site assignment.

### Candidate ORF Identification

The pipeline was implemented in Python (v3.10) using pandas (v1.5), pysam (v0.20), and pybedtools (v0.9). All AUG-initiated ORFs of 30 to 300 codons within annotated transcripts were systematically enumerated genome-wide. This size range was chosen to encompass the canonical sORF definition while excluding trivially short ORFs that are unlikely to encode stable peptides and very long ORFs that overlap with canonical CDS. ORFs overlapping annotated CDS in Ensembl release 109 were excluded, retaining only non-canonical ORFs in 5’ UTRs, 3’ UTRs, lncRNAs, and pseudogene transcripts. Crucially, pseudogene-biotype transcripts from Ensembl were explicitly included in the search space, unlike all three existing benchmarked tools. This stage was applied independently to asynchronous, S phase, and M phase datasets.

### P-site offset calculation

Accurate P-site assignment is essential for evaluating ribosome occupancy and triplet periodicity in Ribo-seq data. RiboWin performs empirical P-site calibration using the psite_calibration.py module. Annotated coding sequences (CDSs) with known reading frames are used to determine optimal offsets for ribosome-protected fragments (RPFs) of different lengths (26–34 nt). For each read length, offsets ranging from 10–16 nucleotides are evaluated, and the offset producing the highest proportion of P-sites in the expected coding frame (Frame 0) is selected. The optimized offsets are stored in a JSON file. The psite_assignment.py module then uses these calibrated offsets to assign P-site positions for all aligned RPF reads. This sample-specific calibration improves the accuracy of downstream translation analyses, including triplet periodicity assessment.

### Frame Periodicity Analysis

For each candidate sORF, P-site reads overlapping all genomic segments were extracted and frame-assigned in a strand-aware manner. Frame 0 fraction was calculated as the proportion of all P-site reads assigned to frame 0. Candidate sORFs were retained if they satisfied two conditions: (i) a minimum of 8 P-site reads across all segments, and (ii) a Frame 0 fraction of at least 0.55. This threshold is substantially above the random expectation of 0.33 for an untranslated sequence and is consistent with established tools including RiboTaper [32]. The choice of 0.55 as a threshold represents a conservative but not overly stringent cutoff, it is sufficient to exclude background noise while remaining below the perfect periodicity values observed for highly expressed canonical CDS, thereby accommodating the lower read depths typical of sORF loci.

### Translation Efficiency Calculation

Translation efficiency (TE) was calculated as the ratio of Ribo-seq P-site read count to RNA-seq read count, a metric widely used to quantify the translational output per unit of transcript [33] . ORFs with zero RNA-seq reads were assigned TE = NaN and excluded from efficiency comparisons. For replicates with RNA-seq read counts below 5, pooled TE was calculated by summing reads across replicates before dividing, to mitigate ratio instability from low count noise. sORFs passed the coverage filter if they had a minimum of 10 Ribo-seq P-site reads in at least one replicate and at least 1 RNA-seq read in at least one replicate. This asymmetric coverage criterion reflects the biological reality that some translated transcripts are present at very low steady-state abundance, as has been observed for certain microprotein-encoding loci where high ribosome occupancy is accompanied by low RNA-seq coverage [30] .

### Dual Biological Replicate Concordance Filter

The defining feature of the pipeline is the requirement that a candidate ORF independently pass all preceding filters in both biological replicates. An ORF was classified as high-confidence only if its orf_id appeared in the high-confidence output of both replicate 1 and replicate 2 for a given condition. This filter was applied independently to the asynchronous, S phase, and M phase datasets. The concordance requirement operationalises the principle that genuine biological translation should be reproducible across independent biological replicates, not merely detectable in a single experiment. This principle is directly analogous to the IDR framework used for ChIP-seq peak calling [34] and the replicate concordance requirements specified by ENCODE for RNA-seq experiments [14] . Unlike Pearson or Spearman correlation-based concordance measures, which allow high-signal outliers to dominate, the binary concordance gate treats each ORF independently and does not allow a highly expressed but single-replicate candidate to pass.

### Benchmarking Against Existing Tools

RiboCode (v1.2.2), Ribo-TISH (v0.2.4), and RibORF (v2.0) were applied to the same asynchronous HeLa Ribo-seq replicates. RiboCode was run with settings -l no -s ATG and default p-value cutoff. Ribo-TISH quality assessment was performed per replicate to determine P-site offsets, followed by ORF prediction with a minimum ORF length of 30 amino acids. For RibORF, P-site offsets were derived from Ribo-TISH quality output and converted to the two-column parameter format, prediction was performed using ribORF.pl with minimum amino acid length 30 and minimum read count 10. The RPL26P19 locus was explicitly queried in all single-replicate and concordant outputs to assess whether each tool was capable of detecting this candidate at any stage of its analysis. This prospective query was registered before examining tool outputs to avoid post-hoc selection bias.

### IGV Validation

High-confidence ORFs were visually validated in the Integrative Genomics Viewer (IGV, v2.16). Single-nucleotide P-site BED files were loaded for all replicates alongside RNA-seq coverage tracks. Assessment criteria included, P-site read concentration within ORF boundaries relative to flanking regions; signal cessation at the stop codon, RNA-seq coverage confirming transcript presence, 3-nucleotide periodicity in the P-site track, and consistency across both biological replicates. IGV validation serves as a qualitative complement to the quantitative computational metrics, enabling visual detection of artefacts such as read pile-ups at splice junctions or regions of high secondary structure that may produce spurious frame periodicity scores.

### Multi-Mapping Assessment

To assess the contribution of multi-mapping reads, alignments were examined using NH tags and mapping quality (MAPQ). Reads with NH: i:1 was considered uniquely mapped. Selected loci were re-evaluated using uniquely mapped reads to determine whether observed ribosome occupancy signals were robust to mapping stringency. This analysis is particularly important for pseudogene loci, which by definition share substantial sequence identity with their parental genes. The retention of concordant signal at RPL26P19 using uniquely mapped reads provides evidence that the observed Ribo-seq signal is not an artefact of reads originating from the highly expressed parental RPL26 locus.

### Database Novelty Verification

Candidate ORF coordinates and sequences were cross-referenced against OpenProt and sORFs.org after pipeline execution and candidate identification, ensuring that pipeline output was generated without prior knowledge of database entries. Where coordinate discrepancies existed between pipeline output and OpenProt entries, amino acid sequence identity was verified by BLASTP. This prospective validation approach eliminates the possibility that database knowledge influenced candidate selection, which is critical for interpreting the significance of the observed concordance between pipeline output and database entries. The post-hoc nature of the database comparison is explicitly stated in the manuscript and is a deliberate methodological choice to preserve the integrity of the validation.

### Multi Cell-Line Application

To assess cell-type specificity of non-canonical translation, the pipeline was applied with full dual replicate concordance to three primary human cell lines (fibroblast, keratinocyte, and melanocyte), each with two biological replicates and matched RNA-seq. Ribo-seq processing followed identical parameters to HeLa. TE was considered calculable only where RNA-seq coverage at the candidate locus was at least 1 read in a given replicate. The inclusion of three distinct normal primary cell types alongside the HeLa cancer cell line enables identification of cell-type-specific translation events and provides an internal specificity control, candidates detected exclusively in cancer cells can be distinguished from those present in normal differentiated cell types.

### Sensitivity analysis

A sensitivity analysis was performed to evaluate the robustness of the translated sORF detection framework under varying filtering thresholds. Two key parameters influencing candidate selection were systematically assessed in HeLa asynchronous (HeLa async) datasets:

**1.** Frame0 fraction threshold (triplet periodicity filter)
**2.** Minimum P-site read count threshold

For the Frame0 fraction analysis, candidate translated ORFs were filtered using progressively stringent thresholds of F0 ≥ 0.50, F0 ≥ 0.55 (current/default threshold), and F0 ≥ 0.60.[32] The number of concordant translated candidates retained across biological replicates was recorded for each condition. Presence of the experimentally relevant candidate RPL26P19 was additionally monitored to assess biological sensitivity. For the minimum read count analysis, translated candidates were filtered using thresholds of ≥ 8, ≥ 10 (current/default threshold), and ≥ 15 P-site reads per replicate. Concordant translated candidates retained after replicate intersection were quantified, and retention/loss of RPL26P19 was examined. This analysis enabled evaluation of the trade-off between detection sensitivity and filtering stringency, while validating the stability of biologically meaningful candidates across parameter settings.

## Results

### Pipeline Design and the Impact of Dual Replicate Concordance on Candidate Numbers

The pipeline was applied to six Ribo-seq datasets (two biological replicates each for asynchronous, S phase, and M phase HeLa cells) along with matched RNA-seq data. The progressive reduction in candidate numbers across pipeline stages illustrates the stringency contribution of each filter. In asynchronous HeLa cells, initial candidate ORF identification yielded 231 and 351 non-canonical ORFs in replicates 1 and 2 respectively. Application of the frame periodicity filter reduced these to 63 and 71 respectively, removing 72.7% of initial candidates and demonstrating that the majority of ribosome-associated non-canonical loci do not show frame-specific translation evidence. The dual-replicate concordance filter produced a further 81.0% reduction, retaining 12 concordant candidates in asynchronous cells. M phase and S phase conditions yielded 9 and 16 concordant candidates respectively.

Across all three conditions, 32 unique sORF loci passed dual-replicate concordance. Of these, 8 were detected exclusively in asynchronous cells, 13 exclusively in S phase, and 7 exclusively in M phase. Two loci were shared between asynchronous and S phase conditions, RPL26P19 was shared between asynchronous and M phase conditions but absent from S phase, and the SMIM15 uORF was present across all three conditions. No sORF was shared exclusively between M phase and S phase. This cell cycle dynamic pattern of non-canonical translation is biologically significant, the fact that only 1 of 32 loci is present across all three conditions indicates that the concordant sORF landscape is actively regulated rather than reflecting constitutive background translation. These results would be entirely obscured by the thousands or millions of low-confidence candidates produced by existing tools, underscoring the analytical value of the stringent concordance approach.

The RiboWin framework generates 632 to 1,192,537-fold fewer concordant candidates than existing tools while being the only approach to detect RPL26P19 consistently across both biological replicates.

The four-tool comparison reveals a striking spectrum of behaviour. RibORF generated 14,780,399 and 19,914,525 candidates in replicates 1 and 2 respectively, with 14,310,443 shared (96.82% concordance). This apparent agreement is artefactual as RibORF effectively calls the vast majority of enumerable ORFs as translated genome-wide, so high concordance simply reflects consistent over-prediction rather than genuine reproducibility. Ribo-TISH presents the opposite failure mode with 842,615 candidates in replicate 1 versus 2,167,749 in replicate 2, a 2.57-fold discrepancy, with only 12.63% shared. The 87.37% of replicate 1 candidates that do not recur in replicate 2 are, by definition, not reproducible translation events, yet all are reported in a standard Ribo-TISH run. RiboCode produced 10,744 and 24,984 candidates with 7,579 shared (70.54% concordance), and is the most informative comparator. RPL26P19 was detected in replicate 2 but absent from replicate 1, meaning it falls into the 29.46% of candidates that are not concordant.

### 3.2 Concordant sORF Landscape Across HeLa Cell Cycle Conditions

#### RPL26P19: Ribo-seq Validation of an Uncharacterised OpenProt AltProt Entry

The ORF at chr5:56,504,634–56,504,933 (minus strand, GRCh38) encodes a predicted 99 amino acid peptide within RPL26P19, an RPL26 pseudogene at the C5orf67 locus on chromosome 5q11.2. Cross-referencing with OpenProt identified a matching AltProt entry at identical coordinates (ENSG00000226221.1) carrying no prior experimental validation no Ribo-seq evidence, no mass spectrometry support, and no unique peptides. The present study provides, to the best of our knowledge, the first experimental computational Ribo-seq evidence for this locus, including dual biological replicate confirmation, quantitative frame periodicity, translation efficiency dynamics, and condition-specific characterisation across asynchronous and M phase cells.

The encoded peptide shares 96.97% amino acid identity with 60S ribosomal protein L26-like 1 (BLASTP E-value: 2e-62, 100% coverage), consistent with RPL26P19’s origin as a processed pseudogene. To address the possibility that the observed signal arises from multi-mapped reads from the highly expressed parental locus, NH tag assessment confirmed that uniquely mapped reads alone support concordant ribosome occupancy at RPL26P19, ruling out a mis-mapping artefact.

Conservation and chromatin evidence further support biological relevance. PhyloP scores indicate nucleotide-level evolutionary constraint across the ORF, and Multiz alignments confirm conservation through birds, amphibians, and teleosts — unexpected for a neutrally evolving pseudogene. ENCODE H3K27Ac marks indicate active chromatin at this locus, and GTEx V8 data confirm RPL26P19 transcription across all 54 profiled human tissues, with notably elevated levels in pancreas (∼15 TPM median). The shared peptide ANGTTVHVGIHPSK was detected in PeptideAtlas (best probability 1.0; 1,374 experiments), including within the smORF-specific proteomics dataset CONTRIB_smORFs_Cui_577993, providing indirect proteomic support. DeepLoc 2.0 predicted cytoplasmic localisation (probability 0.849), consistent with canonical RPL26 biology.

#### Translation Metrics Confirm High-Confidence Status of the RPL26P19 Candidate

Quantitative translation metrics for the RPL26P19 ORF are reported in Table 1. In asynchronous cells, both replicates showed Frame 0 fractions of 0.917 and 0.600 respectively, substantially above the random expectation of 0.33, with a mean TE of 0.731. In M phase cells, Frame 0 fractions of 0.600 and 0.727 were observed with a mean TE of 1.535. Conservative pooled TE estimates were 0.658 for asynchronous and 1.107 for M phase, confirming that the TE increase is not an artefact of low RNA read counts in a single replicate. S phase Ribo-seq detected 8 to 9 reads at this locus per replicate, below the minimum 10-read threshold; TE was therefore not calculated.

**Table 1.** RPL26P19 translation metrics across HeLa cell cycle conditions. TE = translation efficiency (Ribo-seq P-site reads / RNA-seq reads). Frame0 fractions substantially above the random expectation of 0.33 confirm active translation in both asynchronous and M phase. Absence from S phase is consistent across both replicates.

| Stage | Ribo rep1 | Ribo rep2 | RNA rep1 | RNA rep2 | TE rep1 | TE rep2 | TE mean | Frame0 rep1/rep2 |
| --- | --- | --- | --- | --- | --- | --- | --- | --- |
| Asynchronous | 12 | 15 | 26 | 15 | 0.462 | 1.000 | 0.731 | 0.917 / 0.600 |
| S phase | N/A | N/A | N/A | N/A | N/A | N/A | Absent | N/A |
| M phase | 20 | 11 | 23 | 5 | 0.870 | 2.200 | 1.535 | 0.600 / 0.727 |

**Table 2.** Four-tool comparison on asynchronous HeLa Ribo-seq data.

| Feature | RibORF | Ribo-TISH | RiboCode | This framework (RiboWin) |
| --- | --- | --- | --- | --- |
| Rep1 candidates (pre-filter) | 14,780,399 | 842,615 | 10,744 | 231 |
| Rep2 candidates (pre-filter) | 19,914,525 | 2,167,749 | 24,984 | 351 |
| Rep1 ∩ Rep2 concordant | 14,310,443 (96.82%) | 106,429 (12.63%) | 7,579 (70.54%) | 12 (19.05%) |
| Fold more concordant than ours | 1,192,537× | 8,869× | 632× | N/A |
| Novel pseudogene hits rep1 | 0 | 980 | 173 | Explicitly included |
| RPL26P19 detected | Absent both replicates | Absent both replicates | Replicate 2 only; not concordant | Both replicates confirmed |
| Search space | Annotated CDS only | Annotated CDS only | Annotated + some pseudogenes | All biotypes, non-CDS |
| Replicate filter | Not enforced | Not enforced | Not enforced | Mandatory binary |

**Table 3.** Tier 1 Concordant sORF Candidates in Asynchronous HeLa cells. All candidates independently pass frame periodicity and coverage filters in both biological replicates. PC = protein-coding; NMD = nonsense-mediated decay; aa = amino acids; uORF=upstream open reading frame. Frame0 fractions are substantially above the random expectation of 0.33 in all cases.

| sORF ID | Gene | Category | aa | Ribo rep1 | Ribo rep2 | Frame0 rep1 | Frame0 rep2 | Cell cycle conditions |
| --- | --- | --- | --- | --- | --- | --- | --- | --- |
| ORF_13030 | RPL26P19 | Pseudogene | 99 | 12 | 15 | 0.917 | 0.600 | Asynch + M phase |
| ORF_1228 | TMEM218 | Retained intron | 89 | 13 | 212 | 0.923 | 0.769 | Asynch + S phase |
| ORF_4978 | PLEKHG4 | NMD transcript | 93 | 60 | 21 | 0.933 | 0.619 | Asynchronous |
| ORF_13141 | SMIM15 | PC uORF | 86 | 1614 | 5413 | 0.572 | 0.708 | All three conditions |
| ORF_488 | SEPHS1 | PC uORF | 85 | 85 | 23 | 1.000 | 0.913 | Asynchronous |
| ORF_1115 | MEN1 | PC uORF | 98 | 18 | 87 | 1.000 | 0.575 | Asynchronous |
| ORF_2072 | AMIGO2 | PC uORF | 91 | 42 | 49 | 1.000 | 0.612 | Asynchronous |
| ORF_4823 | ADAM10 | PC uORF | 21 | 171 | 47 | 0.626 | 0.830 | Asynchronous |
| ORF_7834 | ZBED6 | PC uORF | 89 | 42 | 162 | 0.952 | 0.574 | Asynchronous |
| ORF_8559 | RERE | PC uORF | 59 | 13 | 88 | 1.000 | 0.716 | Asynchronous |
| ORF_13289 | PPP2R2B | PC uORF | 29 | 25 | 10 | 1.000 | 1.000 | Asynch + S phase |
| ORF_15390 | TMEM65 | PC uORF | 21 | 116 | 13 | 0.552 | 0.846 | Asynchronous |

**Table 4.** Tier 1 Concordant sORF Candidates in HeLa M Phase cells. RPL26P19 is the sole pseudogene-derived candidate, present also in asynchronous cells but absent from S phase. Notable candidates include RUNX1 (implicated in haematopoiesis and leukaemia) and ATXN1 (associated with spinocerebellar ataxia).

| sORF ID | Gene | Category | aa | Ribo rep1 | Ribo rep2 | Frame0 rep1 | Frame0 rep2 | Cell cycle conditions |
| --- | --- | --- | --- | --- | --- | --- | --- | --- |
| ORF_13030 | RPL26P19 | Pseudogene | 99 | 20 | 11 | 0.600 | 0.727 | Asynch + M phase |
| ORF_11751 | HYAL2 | PC uORF | 91 | 65 | 12 | 0.723 | 1.000 | M phase |
| ORF_3960 | ATG2B | PC uORF | 31 | 20 | 15 | 0.700 | 0.800 | M phase |
| ORF_13952 | ATXN1 | PC uORF | 46 | 21 | 12 | 0.667 | 0.917 | M phase |
| ORF_8547 | CDC42BPA | PC uORF | 63 | 69 | 63 | 0.652 | 0.603 | M phase |
| ORF_13141 | SMIM15 | PC uORF | 86 | 600 | 1445 | 0.633 | 0.561 | All three conditions |
| ORF_1618 | FBRSL1 | PC uORF | 88 | 120 | 18 | 0.617 | 0.556 | M phase |
| ORF_8536 | ENAH | PC uORF | 66 | 410 | 143 | 0.590 | 0.650 | M phase |
| ORF_9647 | RUNX1 | PC uORF | 28 | 12 | 61 | 0.583 | 0.738 | M phase |

**Table 5.**
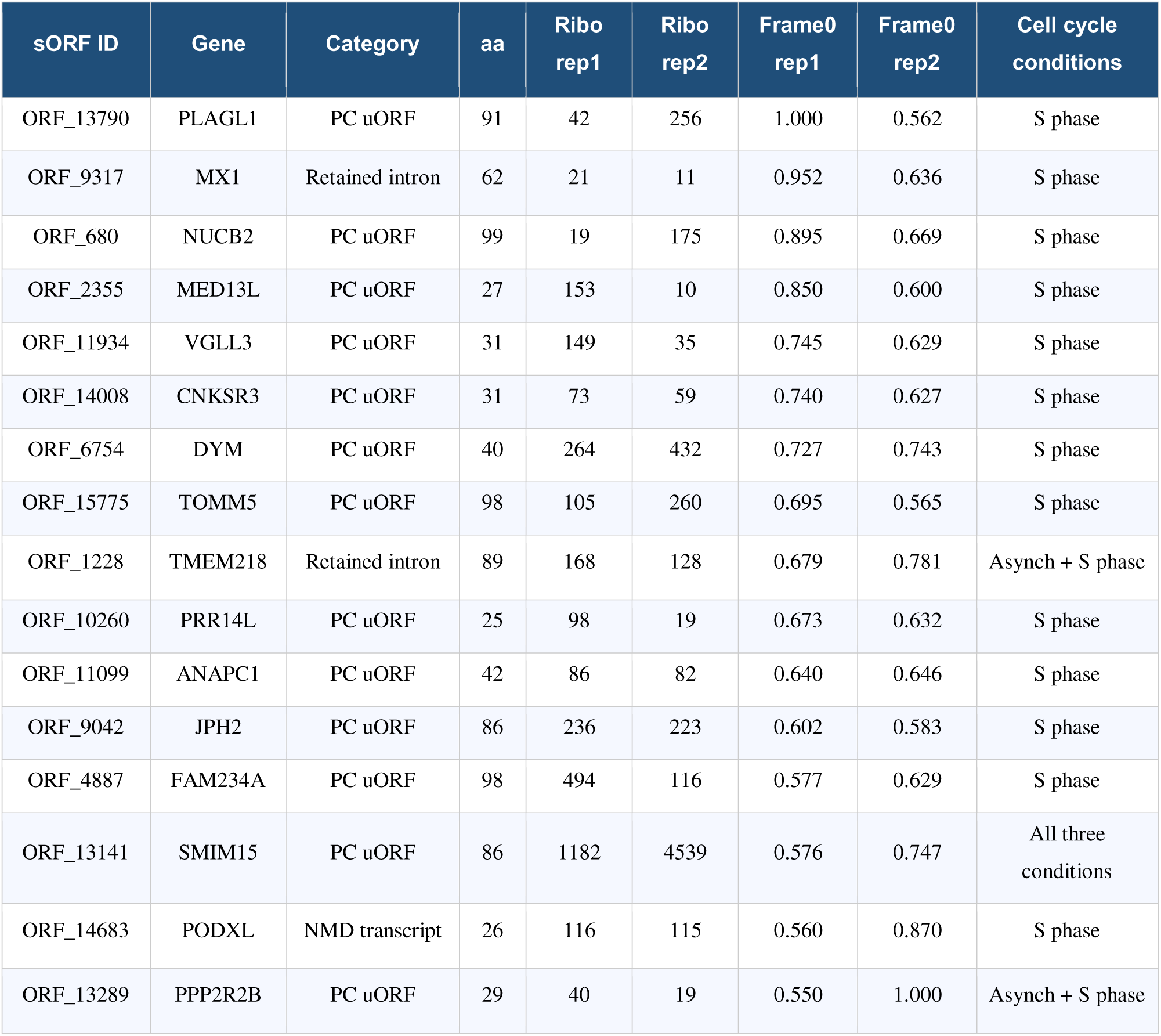
Tier 1 Concordant sORF Candidates in HeLa S Phase cells. 16 candidates are identified, the largest of the three conditions. Notable candidates include ANAPC1 (Anaphase Promoting Complex subunit 1, a core cell cycle regulator), PLAGL1 (an imprinted tumour suppressor), and MX1 (an interferon-stimulated antiviral factor with signal arising from a retained intron).

| sORF ID | Gene | Category | aa | Ribo rep1 | Ribo rep2 | Frame0 rep1 | Frame0 rep2 | Cell cycle conditions |
| --- | --- | --- | --- | --- | --- | --- | --- | --- |
| ORF_13790 | PLAGL1 | PC uORF | 91 | 42 | 256 | 1.000 | 0.562 | S phase |
| ORF_9317 | MX1 | Retained intron | 62 | 21 | 11 | 0.952 | 0.636 | S phase |
| ORF_680 | NUCB2 | PC uORF | 99 | 19 | 175 | 0.895 | 0.669 | S phase |
| ORF_2355 | MED13L | PC uORF | 27 | 153 | 10 | 0.850 | 0.600 | S phase |
| ORF_11934 | VGLL3 | PC uORF | 31 | 149 | 35 | 0.745 | 0.629 | S phase |
| ORF_14008 | CNKSR3 | PC uORF | 31 | 73 | 59 | 0.740 | 0.627 | S phase |
| ORF_6754 | DYM | PC uORF | 40 | 264 | 432 | 0.727 | 0.743 | S phase |
| ORF_15775 | TOMM5 | PC uORF | 98 | 105 | 260 | 0.695 | 0.565 | S phase |
| ORF_1228 | TMEM218 | Retained intron | 89 | 168 | 128 | 0.679 | 0.781 | Asynch + S phase |
| ORF_10260 | PRR14L | PC uORF | 25 | 98 | 19 | 0.673 | 0.632 | S phase |
| ORF_11099 | ANAPC1 | PC uORF | 42 | 86 | 82 | 0.640 | 0.646 | S phase |
| ORF_9042 | JPH2 | PC uORF | 86 | 236 | 223 | 0.602 | 0.583 | S phase |
| ORF_4887 | FAM234A | PC uORF | 98 | 494 | 116 | 0.577 | 0.629 | S phase |
| ORF_13141 | SMIM15 | PC uORF | 86 | 1182 | 4539 | 0.576 | 0.747 | All three conditions |
| ORF_14683 | PODXL | NMD transcript | 26 | 116 | 115 | 0.560 | 0.870 | S phase |
| ORF_13289 | PPP2R2B | PC uORF | 29 | 40 | 19 | 0.550 | 1.000 | Asynch + S phase |

**Table 6.** UBE3AP2 (sORF 9495) translation metrics across human cell lines. N/C = not calculable due to zero RNA-seq coverage. The reciprocal specificity pattern UBE3AP2 present in normal cell types, absent in HeLa; RPL26P19 present in HeLa, absent in normal cell types suggests active, cell-type specific regulation of pseudogene-derived translation.

| Cell line | Type | Ribo rep1 | Ribo rep2 | RNA rep1 | RNA rep2 | TE rep1 | TE rep2 | Frame0 rep1/rep2 | Status |
| --- | --- | --- | --- | --- | --- | --- | --- | --- | --- |
| HeLa | Cancer | N/A | N/A | N/A | N/A | N/A | N/A | N/A | Absent |
| Fibroblast | Normal | N/A | N/A | N/A | N/A | N/A | N/A | 0.26 / 0.26 | Sub-threshold F0 |
| Keratinocyte | Normal | 37 | 85 | 0 | 0 | N/C | N/C | 0.89 / 0.97 | Tier 1 |
| Melanocyte | Normal | 77 | 93 | 6 | 0 | 12.833 | N/C | 0.89 / 0.90 | Tier 1 |

TE increases 2.1-fold from asynchronous cells (0.731) to M phase (1.535), consistent with translational upregulation during mitosis. The elevated TE_rep2 in M phase (2.200) reflects low RNA-seq coverage in replicate 2 (5 reads); the pooled TE of 1.107 confirms that the directional increase is genuine. The observed anti-correlation between RNA levels and translation efficiency across cell cycle conditions is consistent with post-transcriptional regulation and is a hallmark of translationally regulated genes [1] ; [33] . The finding that total Ribo-seq reads at the locus increase from 27 to 31 while RNA reads decrease from 41 to 28 between asynchronous and M phase conditions argues strongly against the possibility that the TE increase is driven by RNA-seq read count artefacts.

#### Cell Cycle Translation Pattern of RPL26P19 as a Biological Specificity Control

The translation pattern at RPL26P19 (concordant signal in asynchronous and M phase replicates, consistently absent from both S phase replicates) is not consistent with random noise or technical meaningful oscillatory pattern across independent datasets. The quantitative increase in TE in M phase occurs against a backdrop of reduced RNA levels, with total RNA reads decreasing from 41 in asynchronous conditions to 28 in M phase while Ribo-seq reads increased from 27 to 31. This indicates that the signal reflects genuine changes in translational activity rather than transcript abundance.

The observed pattern is consistent with cap-independent translation during mitosis. [27] demonstrated that cap-dependent translation is broadly suppressed during M phase, while certain mRNAs that contain internal ribosome entry sites (IRES) or other cap-independent regulatory elements escape this suppression and may be translationally upregulated. The elevated TE of RPL26P19 in M phase is therefore consistent with a cap independent mechanism, which would be consistent with the retention of translational regulatory elements from the parental RPL26 locus. [28] characterised cell cycle regulated IRES activity in several mRNAs, providing a mechanistic precedent for the type of translational regulation observed here.

The S phase absence provides the pipeline with a biologically grounded internal specificity control independent of any external knowledge about the locus. The fact that sub threshold signal of 8 to 9 reads per replicate is detected in S phase but does not meet the minimum 10 read filter is noteworthy because it suggests low level rather than absent translation in S phase, consistent with cell cycle regulated modulation of translation efficiency rather than complete transcriptional silencing of the locus during DNA replication.

#### IGV Validation Confirms P-site Periodicity and ORF Boundary Specificity

Visual inspection of P-site tracks in IGV confirmed all computational metrics at nucleotide resolution. In both asynchronous replicates, P-site reads were concentrated within the RPL26P19 ORF boundaries (chr5:56,504,635 to 56,504,934) with substantially lower read density in flanking intronic regions. Signal cessation at the predicted stop codon was visible in both replicates. In M phase, both replicates showed increased P-site read density within the ORF relative to asynchronous cells, qualitatively consistent with the elevated TE values. The 3-nucleotide periodicity characteristic of active elongating ribosomes was visually apparent. S phase tracks for both replicates showed an absence of concentrated P-site signal at the locus, consistent with sub-threshold read counts. The concordance between IGV visual inspection and the computational metrics across all six Ribo-seq datasets (two replicates per condition across three conditions) provides strong independent validation of the automated pipeline results.

#### Multi Cell Line Analysis Reveals a Cancer versus Normal Dichotomy in Pseudogene-Derived Translation

The framework was applied with full dual-replicate concordance to three primary human cell lines (fibroblast, keratinocyte, and melanocyte) alongside HeLa, enabling a direct cancer versus normal comparison. RPL26P19 failed to pass dual-replicate concordance in any of the three normal primary cell lines. In fibroblast, both replicates showed Frame0 fractions of 0.26, below the 0.55 threshold. In keratinocyte and melanocyte, RPL26P19 was absent from concordant output entirely. This pattern indicates that RPL26P19 translation is cancer cell-specific under stringent criteria.

Conversely, sORF 9495 within UBE3AP2 (chr21:31,062,633 to 31,062,911, 92 amino acids), a pseudogene of the E3 ubiquitin ligase UBE3A, passed dual replicate concordance in keratinocyte and melanocyte but was absent from HeLa. Frame0 fractions were consistently high (keratinocyte replicate 1: 0.89, replicate 2: 0.97; melanocyte replicate 1: 0.89, replicate 2: 0.90). Translation efficiency for UBE3AP2 could be calculated in melanocyte only, where one replicate had detectable RNA-seq coverage (6 reads), yielding TE_rep1 = 12.833. This pattern of high Ribo-seq signal with low or absent RNA-seq coverage is consistent with efficient translation from low abundance transcripts, as has been observed in certain microprotein-encoding loci (Huang et al., 2020).

#### Prospective Validation: UBE3AP2 Ribo-seq Signal Corroborates a Proteomics-Confirmed OpenProt Entry

Cross-referencing UBE3AP2 sORF_9495 with OpenProt revealed AltProt entry ENSG00000228941.1 (92 amino acids, 10.79 kDa, chr21:31,061,820–31,062,098). The coordinate offset (∼813 bp) relative to the predicted sORF reflects differing UBE3AP2 transcript isoform models; BLASTP confirmed identical amino acid sequences. The OpenProt entry is supported by 3 PSMs from 2 unique tryptic peptides [24] , providing independent proteomics corroboration.

This constitutes prospective validation, UBE3AP2 sORF_9495 was identified from Ribo-seq signal alone across three independent normal cell lines, with the proteomics confirmation discovered post-hoc. By contrast, RPL26P19 corresponds to an OpenProt AltProt entry at identical coordinates with no prior experimental support, making this study, to the best of our knowledge, the first experimental computational Ribo-seq validation of that computational prediction. Together, the two candidates delineate complementary validation outcomes UBE3AP2 demonstrates Ribo-seq independently corroborating proteomics, while RPL26P19 demonstrates Ribo-seq providing primary experimental evidence where proteomics has not yet reached. This complementarity strengthens the interpretation that both loci represent genuinely translated candidates rather than false positives.

The prospective design is methodologically significant. By completing candidate identification before any database cross-referencing, and documenting the order of operations explicitly, the study demonstrates that candidate selection is driven by Ribo-seq signal rather than pre-existing database knowledge an important safeguard against the implicit confirmation bias that arises when database searches follow, rather than precede, candidate reporting.

#### Sensitivity Analysis: Frame0 and Minimum Read Count Threshold

Sensitivity analysis demonstrated that translation candidate recovery was strongly influenced by filtering thresholds. In the Frame0 fraction threshold analysis (Figure 6A), relaxing the periodicity threshold to F0 ≥ 0.50 increased the number of concordant translated candidates from 12 to 20, whereas increasing stringency to F0 ≥ 0.60 reduced the candidate count to 8. Importantly, the candidate RPL26P19 remained consistently detected across all tested Frame0 thresholds, indicating stable translational periodicity and robustness to periodicity filtering parameters. In the minimum P-site read count analysis (Figure 5B), the default threshold of ≥ 10 reads retained 12 concordant candidates, while a more lenient threshold (≥ 8 reads) increased candidate recovery slightly to 13. However, increasing the threshold to ≥ 15 reads reduced the number of candidates to 7 and resulted in the loss of RPL26P19, as replicate-level ribosome occupancy for the candidate remained below the stricter cutoff (12 and 15 reads per replicate). Overall, the analysis demonstrated that the selected default thresholds (F0 ≥ 0.55 and ≥ 10 P-site reads) provided an effective balance between sensitivity and specificity, preserving biologically relevant translated candidates while limiting low-confidence detections.

**Figure 1.**
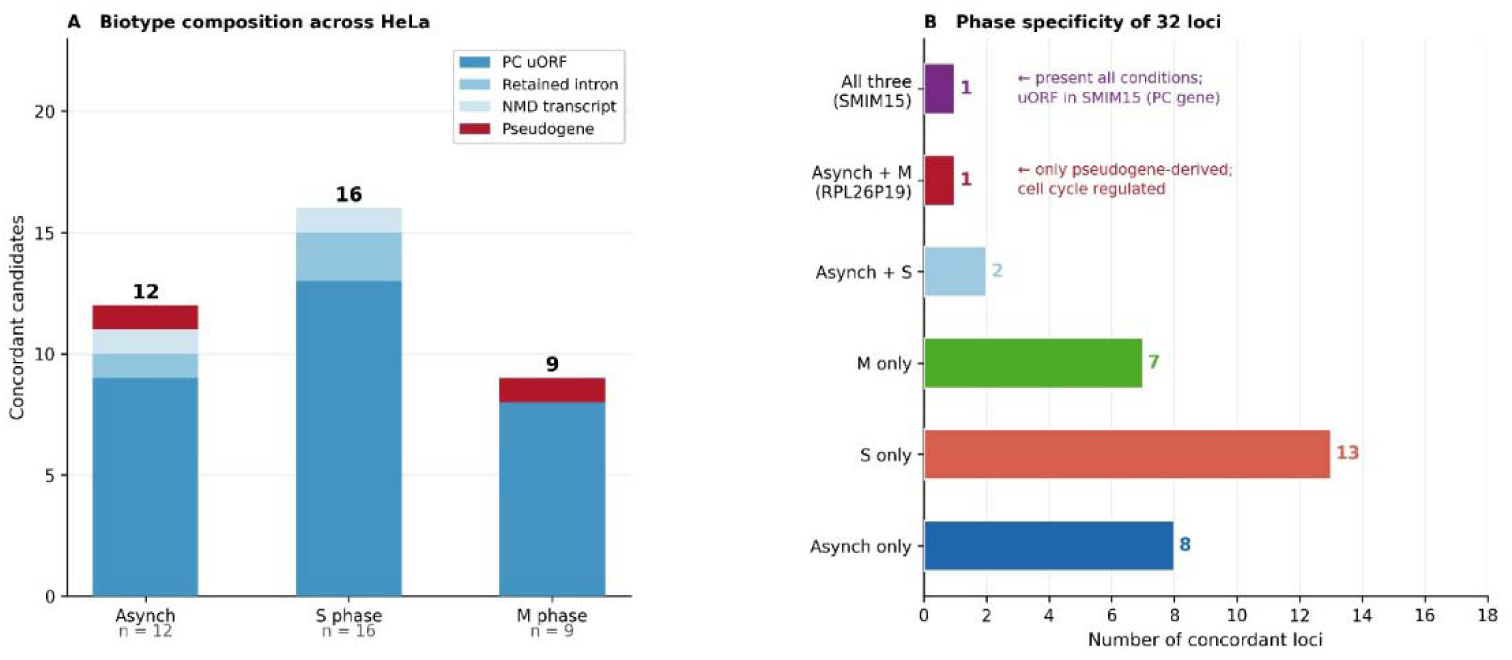
Non-canonical translatome landscape across HeLa cell cycle stages. (A) Biotype composition of concordant sORF candidates per condition; RPL26P19 (pseudogene, red) is annotated. (B) Phase specificity of 32 concordant loci, quantifying loci unique to each condition or shared across conditions.

**Figure 2.**
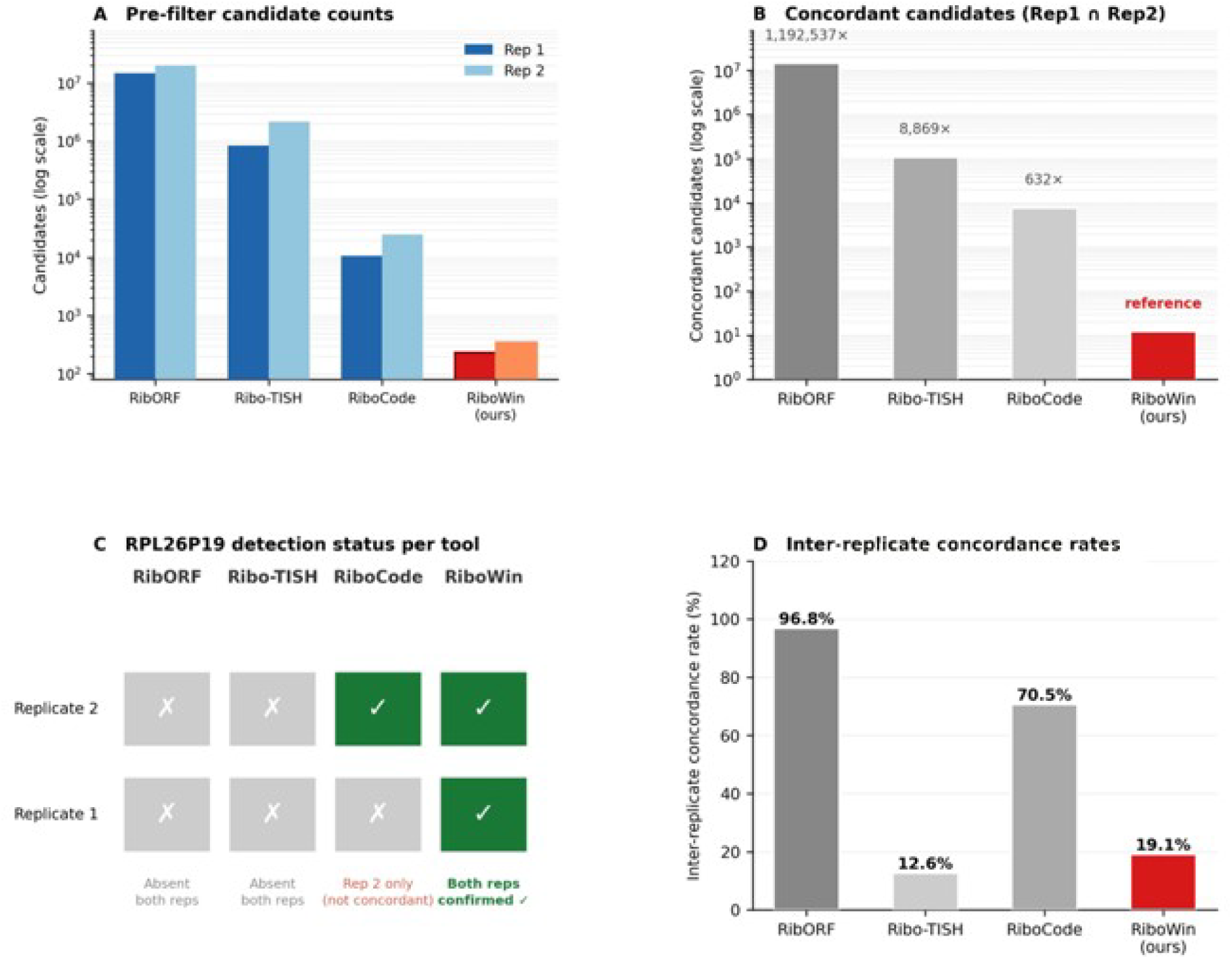
Four-tool comparison on asynchronous HeLa Ribo-seq data. (A) Pre-filter candidate counts per tool across two biological replicates on a log scale. (B) Concordant candidates (Rep1 ∩ Rep2) for each tool, with fold-excess over RiboWin annotated. (C) RPL26P19 detection status per tool across replicates. (D) Inter-replicate concordance rates; note that high concordance in RibORF reflects systematic over-prediction rather than specificity.

**Figure 3:**
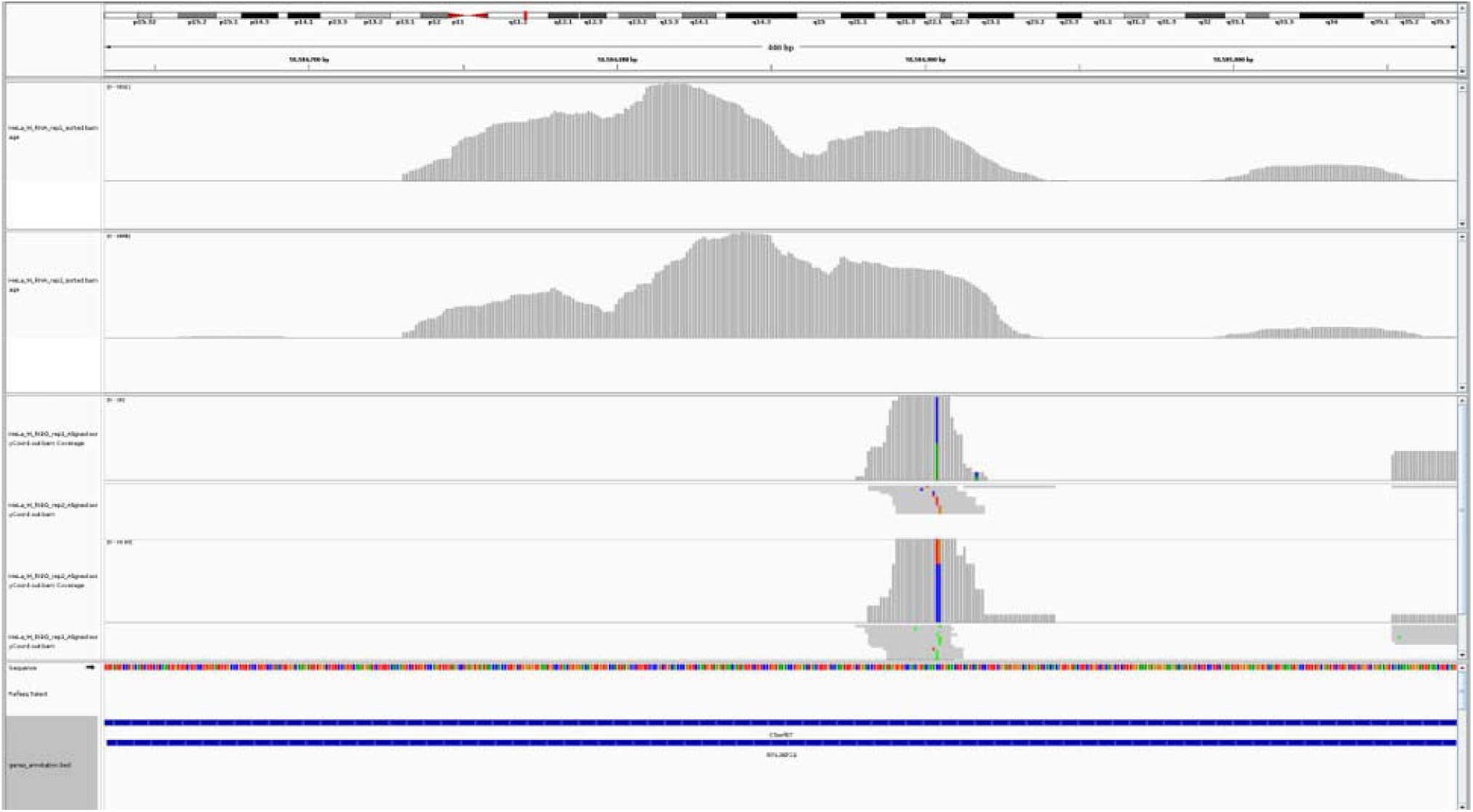
**IGV Validation for M phase.**

**Figure 4:**
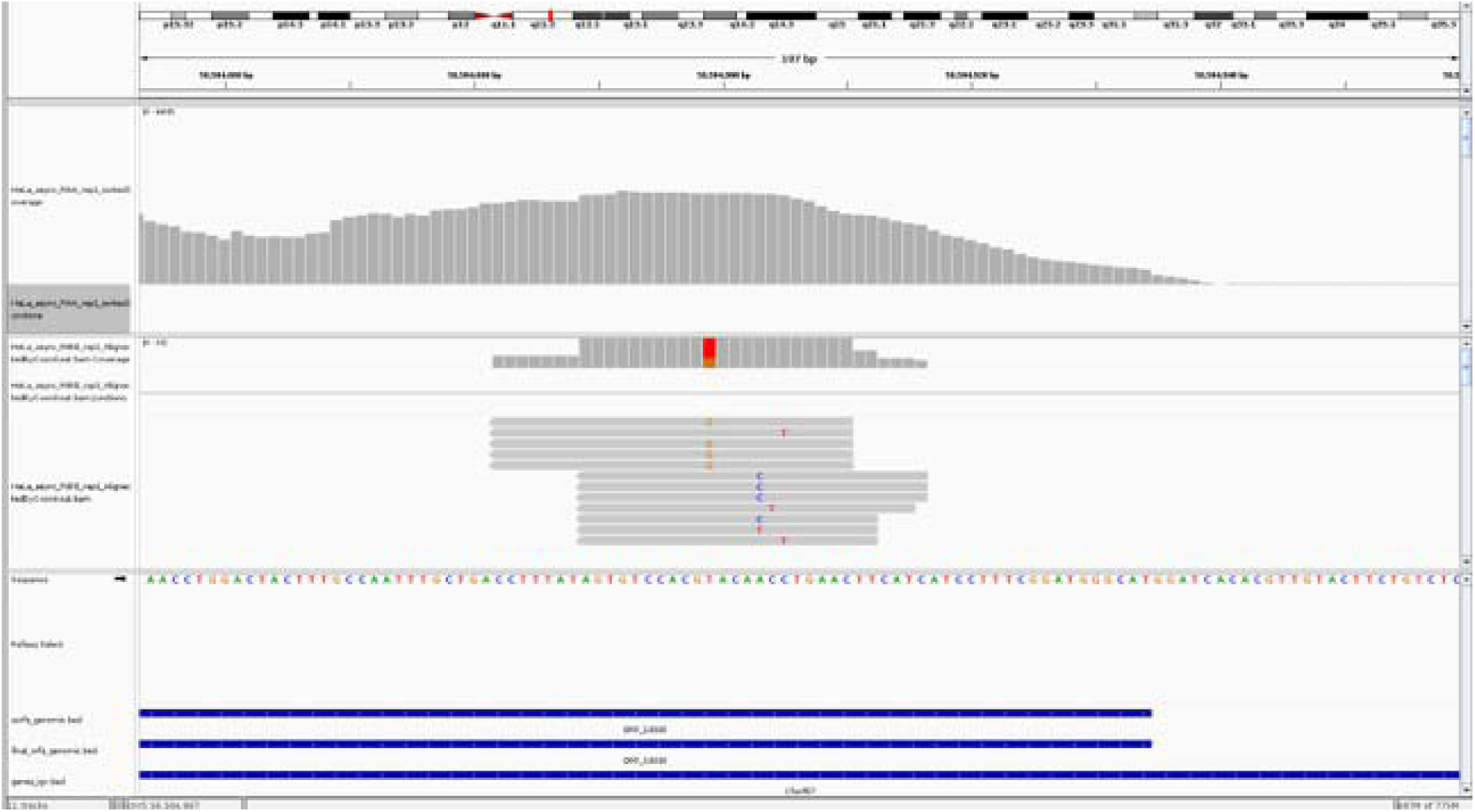
**IGV Validation for Asynchronous stage.**

**Figure 5.**
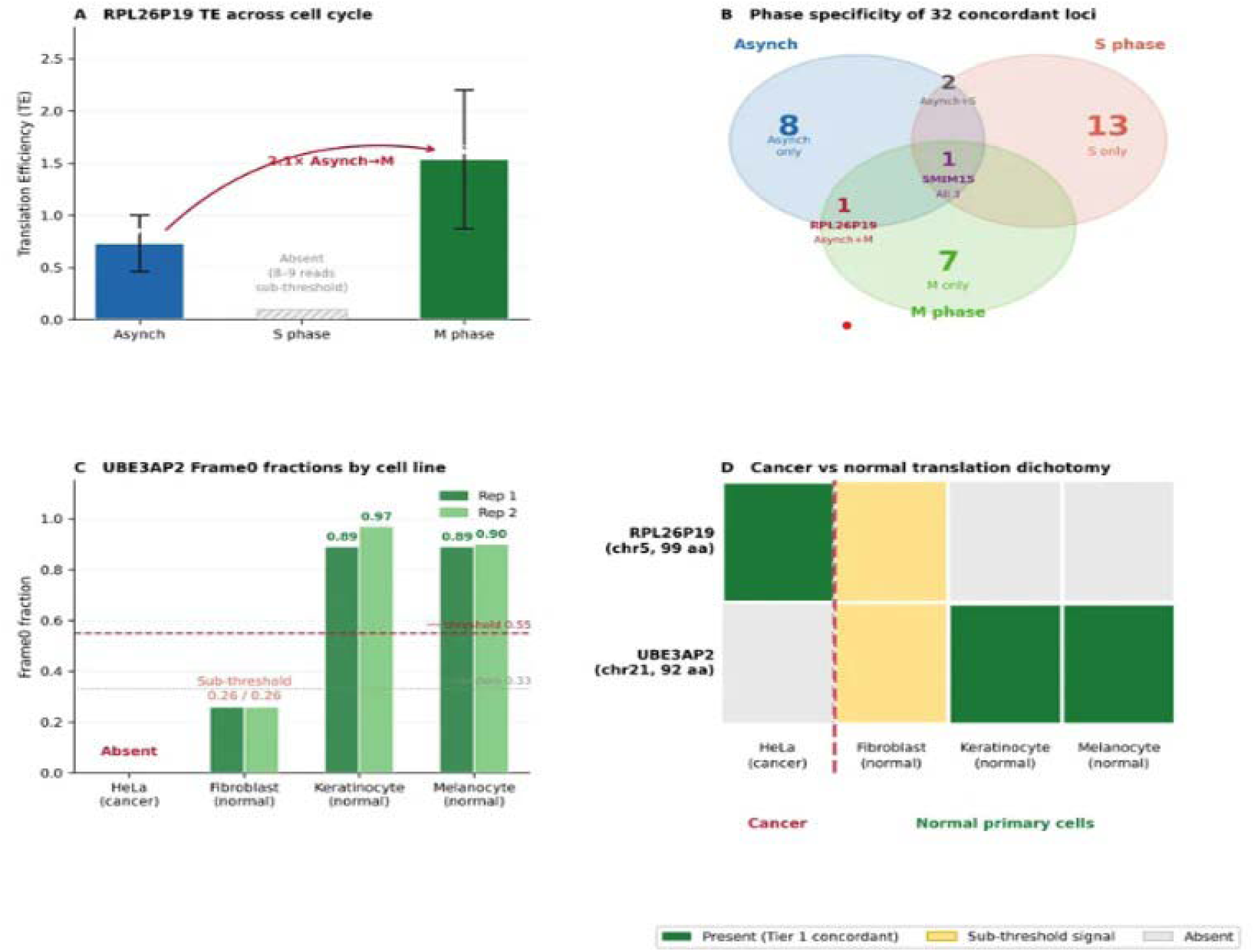
RPL26P19 and UBE3AP2: cell cycle dynamics and cancer/normal specificity. (A) RPL26P19 translation efficiency (TE) across cell cycle conditions; S phase signal is sub-threshold. (B) Phase specificity Venn diagram for 32 concordant loci across asynchronous, S phase, and M phase conditions. (C) UBE3AP2 Frame0 fractions by cell line showing concordant Tier 1 signal in keratinocyte and melanocyte but absence in HeLa. (D) Cancer versus normal translation dichotomy for RPL26P19 and UBE3AP2 across the four cell lines.

**Figure 6:**
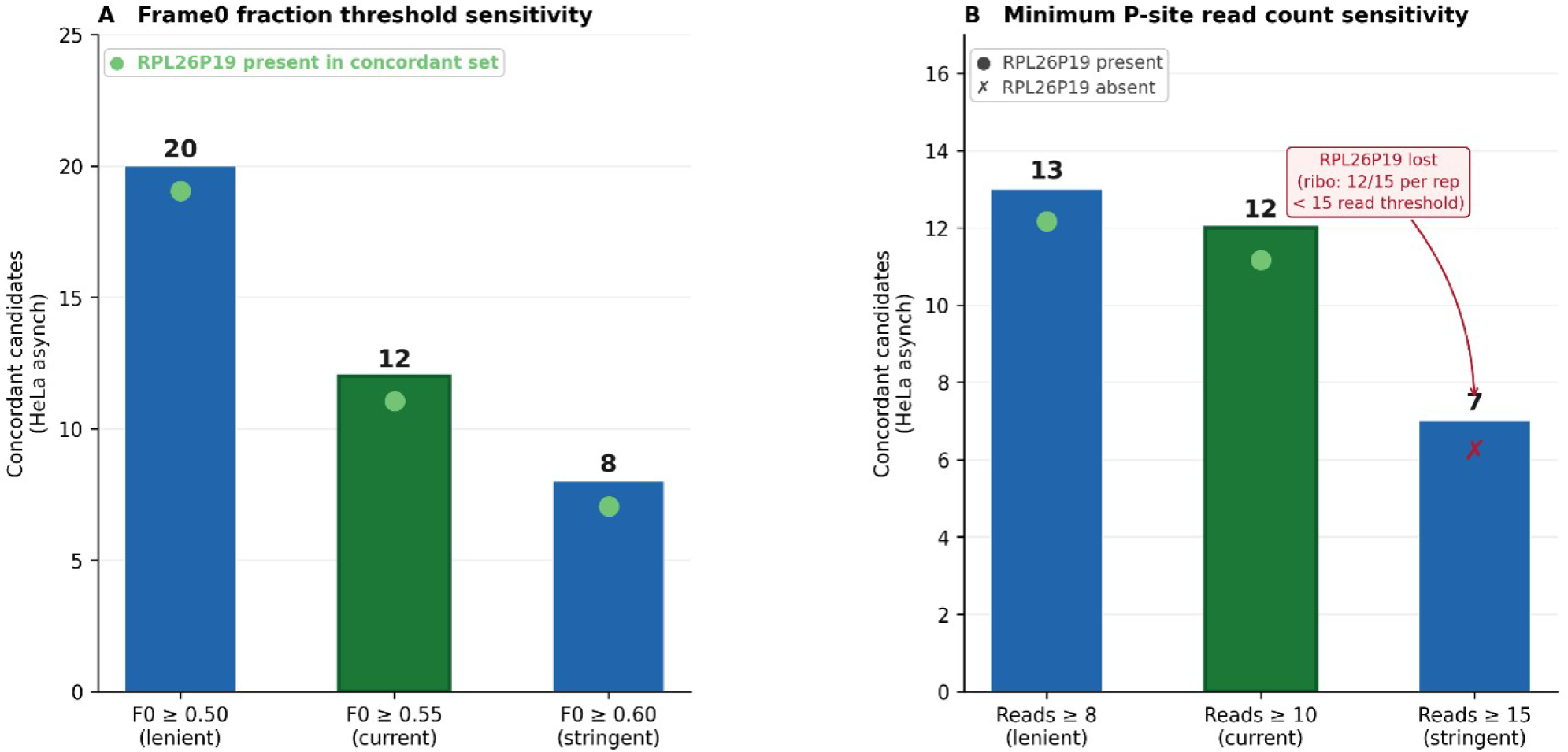
Sensitivity Analysis. (A) Frame0 threshold sensitivity check for 0.50, 0.55, and 0.60. (B) Minimum P-site read count sensitivity for reads greater than 8, 10, and 15.

## Discussion

### Pseudogene inclusive search space and replicate concordance as complementary contributions

The pipeline makes two distinct and complementary contributions to the sORF discovery field. The first is a limited search space of non-canonical that explicitly includes pseudogene transcripts, a class systematically excluded from all three existing tools. The second is dual biological replicate concordance as a mandatory binary filter applied at the candidate selection stage rather than used solely for downstream quantification. Together, these two design choices explain why the pipeline recovers RPL26P19 while RiboCode, Ribo-TISH, and RibORF do not.

The four-tool comparison reveals a striking spectrum of behaviour. RibORF generated 14,780,399 and 19,914,525 candidates in replicates 1 and 2 respectively, with 14,310,443 shared (96.82% concordance). This apparent agreement is artefactual as RibORF effectively calls the vast majority of enumerable ORFs as translated genome-wide, so high concordance simply reflects consistent over-prediction rather than genuine reproducibility. The RPL26P19 locus was absent from all RibORF output, demonstrating that an inflated search space without quality filtering does not resolve the pseudogene detection problem. Ribo-TISH presents the opposite failure mode which is 842,615 candidates in replicate 1 versus 2,167,749 in replicate 2, a 2.57-fold discrepancy, with only 12.63% shared. The 87.37% of replicate 1 candidates that do not recur in replicate 2 are, by definition, not reproducible translation events, yet all are reported in a standard Ribo-TISH run. RPL26P19 was absent from Ribo-TISH output in both replicates, likely because Ribo-TISH restricts its search to annotated coding regions and therefore cannot interrogate the pseudogene locus at all. RiboCode produced 10,744 and 24,984 candidates with 7,579 shared (70.54% concordance), and is the most informative comparator. RPL26P19 was detected in replicate 2 but absent from replicate 1, meaning it falls into the 29.46% of candidates that are not concordant and would therefore not be reported. The signal is real, RiboCode is sensitive enough to detect it but without a concordance requirement, there is no mechanism to distinguish this genuine event from the thousands of other single-replicate detections. The pipeline resolves this by treating concordance as a hard binary gate. Applied to the same data, it identifies 12 candidates from asynchronous HeLa cells, 632-fold fewer than RiboCode, 8,869-fold fewer than Ribo-TISH, and 1,192,537-fold fewer than RibORF, and is the only approach to report RPL26P19 consistently across both replicates. The requirement for dual replicate concordance is already standard in ChIP-seq [14] ; [15] and virtually every other quantitative genomics assay; applying the same principle to sORF discovery does not represent an arbitrary stringency increase but an alignment with reproducibility standards that the field has otherwise adopted.

### RPL26P19 as a proactive case of the pipeline’s evidentiary framework

RPL26P19 provides a representative example of the biological value of extending sORF interrogation beyond conventional protein-coding annotations. Although the locus is annotated as a processed pseudogene and lacks an annotated canonical coding sequence, RiboWin consistently identified an ORF at this locus across independent biological replicates, with P-site occupancy concentrated within the predicted ORF and frame-0 fractions of 0.917 and 0.600 in asynchronous replicates and 0.600 and 0.727 in M-phase replicates. The corresponding IGV profiles further demonstrated ORF-restricted P-site accumulation, signal termination near the predicted stop codon, and three-nucleotide periodicity, providing nucleotide-level support for the computational classification. Importantly, independent transcriptional evidence further supports activity at this locus. Examination of FANTOM5 Cap Analysis of Gene Expression (CAGE) data identified a CAGE-supported transcription start site (TSS) at chr5:56,505,108–56,505,118 (GRCh38), overlapping the 5′ end of the RPL26P19 transcript [35] . Given the minus-strand orientation of RPL26P19, the region toward increasing genomic coordinates represents the transcriptional upstream direction; consequently, the region upstream of this TSS can be considered a putative promoter region. Together with the surrounding ENCODE regulatory annotations examined at the locus, the FANTOM5 CAGE signal provides orthogonal evidence that RPL26P19 is an actively transcribed genomic region, complementing the ribosome-profiling evidence obtained by RiboWin. The significance of RPL26P19 in this study therefore lies not in the discovery of a previously unknown pseudogene, but in demonstrating the ability of the RiboWin framework to interrogate annotated non-canonical transcript classes while retaining stringent evidence for candidate selection. The combination of a pseudogene inclusive search space with mandatory dual-replicate concordance distinguishes RiboWin from approaches in which candidate detection is primarily driven by individual replicate level signals. In the benchmarking analysis, RPL26P19 was absent from RibORF and Ribo-TISH, whereas RiboCode detected the locus in only one replicate and consequently did not retain it as a concordant candidate. RiboWin, in contrast, recovered the locus in both biological replicates. Thus, the RPL26P19 example illustrates the central methodological principle of RiboWin which is the expansion of the search space alone is insufficient unless accompanied by stringent reproducibility criteria. The independent FANTOM5 and examination from UCSC Genome Browser [36] supported TSS and inferred promoter context provide additional biological support for the transcriptional activity of the locus, while the reproducible P-site signal and frame periodicity provide evidence at the translational level. Collectively, these observations strengthen the interpretation of RPL26P19 as a biologically credible non-canonical sORF candidate and demonstrate how RiboWin can reduce a broad search space to a small set of reproducibly supported candidates.

### Why Ribo-seq rather than RNA-seq alone

The detection of RPL26P19 and UBE3AP2 as translated loci requires ribosome profiling data and could not have been achieved from RNA-seq alone. RNA-seq measures transcript abundance, not ribosome occupancy, and provides no information about whether a given ORF is being actively decoded. Non-coding transcripts, pseudogene-derived RNAs, and translated sORF-containing transcripts are all transcribed and would appear in an RNA-seq dataset with equal or greater read depth than in Ribo-seq, yet transcript presence does not imply translation [12] . For small ORFs in particular, the translated peptide is too short to be predicted from sequence context alone with acceptable specificity, and the reading frame information embedded in Ribo-seq P-site periodicity is the primary evidence of active elongation. In the case of UBE3AP2, both keratinocyte replicates had zero RNA-seq reads at the locus, meaning RNA-seq would have excluded this candidate entirely, while Ribo-seq detected robust concordant signal with Frame 0 fractions of 0.89 and 0.97. RNA-seq also lacks the nucleotide resolution necessary to assign reads to a specific frame, which is the defining metric used here to discriminate translating from scanning or stalled ribosomes [5] . Ribo-seq is therefore not merely complementary to RNA-seq for this task; it is the prerequisite technology for detecting translated sORFs in non-canonical loci.

### Prospective validation and the significance of post-hoc OpenProt corroboration

Both candidates were independently corroborated by OpenProt entries discovered only after pipeline execution, providing prospective rather than confirmatory validation. For RPL26P19, OpenProt lists an AltProt entry at identical coordinates with no experimental support of any kind no Ribo-seq evidence, no mass spectrometry, and no unique peptides. The Ribo-seq evidence presented here constitutes, to our knowledge, the first experimental support for this entry, and the condition specific characterisation absent in S phase and 2.1-fold elevated in translation efficiency during M phase represents a level of biological resolution absent from any database annotation. For UBE3AP2, the corresponding OpenProt entry carries 3 PSMs from 2 unique tryptic peptides [24] ; the pipeline independently identifies the same protein from Ribo-seq signal across three normal primary cell lines without any prior awareness of this mass spectrometry support. The two candidates therefore occupy complementary positions in the validation landscape, RPL26P19 is a Ribo-seq validated but proteomically unconfirmed entry, while UBE3AP2 demonstrates independent convergence between ribosome profiling and proteomics. The absence of prior knowledge of these database entries during analysis is important as it eliminates confirmation bias as an explanation for the results and demonstrates that the pipeline’s candidate selection is driven by the Ribo-seq signal rather than by database annotation

### Cell cycle staging as a pipeline specificity control

The use of matched asynchronous, S phase, and M phase datasets provides an internal specificity control that is independent of any external database knowledge. The oscillatory translation pattern at RPL26P19 concordant signal in asynchronous and M phase replicates, consistently absent from both S phase replicates is not consistent with random ribosomal noise, which would not be expected to replicate a biologically coherent cell-cycle pattern across independent datasets. The 2.1-fold increase in translation efficiency from asynchronous to M phase occurs against a backdrop of reduced RNA-seq counts, with total Ribo-seq reads increasing from 27 to 31 while RNA reads fell from 41 to 28. This anti-correlation is consistent with translational upregulation during mitosis when cap-dependent translation is broadly suppressed [27] , suggesting that RPL26P19 translation may be driven by a cap-independent mechanism in M phase. The S phase absence is consistent with stage-specific translational repression during the DNA replication window and provides the pipeline with a biologically grounded specificity readout that existing tools, which do not enforce concordance across conditions, cannot exploit. Of the 32 unique concordant loci identified across all three conditions, only one the SMIM15 uORF with 5.95 TE_mean in asynchronous, 2.09 in M phase and 5.35 in S phase is present across all three, while 23 of 32 are condition specific, demonstrating that the concordant sORF landscape is dynamically regulated rather than static an observation that would be obscured by the thousands of low-confidence candidates reported by existing tools.

## Conclusion

This study presents RiboWin, a non-canonical sORF discovery pipeline incorporating a pseudogene-inclusive search space and mandatory dual biological replicate concordance, applied to four human cell lines with matched biological replicates. The four-tool comparison against RiboCode, Ribo-TISH, and RibORF demonstrates a concordance rate of 19.1%, orders of magnitude more stringent than all existing tools, while recovering pseudogene-derived candidates invisible to all three benchmarked approaches. Across asynchronous, S phase, and M phase HeLa conditions, 32 unique concordant sORF loci were identified spanning protein-coding uORFs, retained introns, NMD transcripts, and one pseudogene-derived candidate (RPL26P19). The cell-cycle-dynamic pattern of this landscape, with only 1 of 32 loci present across all three conditions, demonstrates that the reproducibly translated non-canonical proteome is actively regulated in a phase-specific manner.

Two candidates survive dual replicate concordance across four human cell lines. RPL26P19 provides, to the best of our knowledge, the first Ribo-seq-based experimental evidence supporting an unvalidated OpenProt prediction, with condition-specific characterisation showing increased translation during M phase relative to asynchronous cells and absence in S phase. UBE3AP2 independently supports a proteomics-based OpenProt annotation across three primary human cell types and is not detected in the HeLa dataset. The observed difference between HeLa and the primary cell types analysed suggests that pseudogene derived non-canonical translation is regulated in a cell type specific manner, with one candidate exhibiting cancer-specific translation and another exhibiting normal-tissue specificity.

Together, these findings indicate that pseudogene loci represent a previously underexplored source of translated sORFs that existing pipelines do not systematically capture. The methodological advances introduced by RiboWin particularly the pseudogene-inclusive search space and the mandatory binary concordance filter establish a framework that can be applied to any Ribo-seq dataset with two or more biological replicates, and that can be extended to additional non-canonical transcript classes as genome annotation databases continue to expand. The broader message is clear that the non-canonical translatome is richer, more regulated, and more biologically meaningful than current tools reveal. Addressing the twin limitations of restricted search space and absent concordance requirements is essential for the field to move from large catalogues of unvalidated predictions to small, high-confidence sets of biologically interpretable candidates.

## Supporting information

supp doc.

## Acknowledgement

Department of Biotechnology, M. S. Ramaiah Institute of Technology, MSR Nagar, Bengaluru, India

Author Contributions Statement:

VD, AN, BYK, and YSR contributed to the conception and design of the experiments, experimental investigation, data analysis, interpretation of the results, and drafting of the manuscript. SP and BV contributed to data analysis, interpretation of the results, and drafting of the manuscript. All authors reviewed and approved the final version of the manuscript for submission.

## Data availability

All data used to support the findings of this study are included within the article.

## Conflict of interest

The authors declare that they have no conflicts of interest with the contents of this article.

## Notes

### Competing Interest Statement

The authors have declared no competing interest.

## References

1. Ingolia NT, Ghaemmaghami S, Newman JRS, Weissman JS. Genome-wide analysis in vivo of translation with nucleotide resolution using ribosome profiling. Science (1979). 2009 Apr 10;324(5924):218–23. doi:10.1126/SCIENCE.1168978;PAGEGROUP:STRING:PUBLICATION PubMed PMID: 19213877.

2. Couso JP, Patraquim P. Classification and function of small open reading frames. Nature Reviews Molecular Cell Biology 2017 18:9. 2017 Jul 12;18(9):575–89. doi:10.1038/nrm.2017.58 PubMed PMID: 28698598.

3. Brar GA, Weissman JS. Ribosome profiling reveals the what, when, where and how of protein synthesis. Nature Reviews Molecular Cell Biology 2015 16:11. 2015 Oct 14;16(11):651–64. doi:10.1038/nrm4069 PubMed PMID: 26465719.

4. Anderson DM, Anderson KM, Chang CL, Makarewich CA, Nelson BR, McAnally JR, Kasaragod P, Shelton JM, Liou J, Bassel-Duby R, Olson EN. A micropeptide encoded by a putative long noncoding RNA regulates muscle performance. Cell. 2015 Feb 12;160(4):595–606. doi:10.1016/j.cell.2015.01.009 PubMed PMID: 25640239.

5. Ingolia NT, Brar GA, Stern-Ginossar N, Harris MS, Talhouarne GJS, Jackson SE, Wills MR, Weissman JS. Ribosome Profiling Reveals Pervasive Translation Outside of Annotated Protein-Coding Genes. Cell Rep. 2014 Sep 11;8(5):1365–79. doi:10.1016/j.celrep.2014.07.045 PubMed PMID: 25159147.

6. Ji Z, Song R, Regev A, Struhl K. Many lncRNAs, 5′UTRs, and pseudogenes are translated and some are likely to express functional proteins. Elife. 2015 Dec 19;4(DECEMBER2015). doi:10.7554/ELIFE.08890 PubMed PMID: 26687005.

7. Bazzini AA, Johnstone TG, Christiano R, MacKowiak SD, Obermayer B, Fleming ES, Vejnar CE, Lee MT, Rajewsky N, Walther TC, Giraldez AJ. Identification of small ORFs in vertebrates using ribosome footprinting and evolutionary conservation. EMBO Journal. 2014 May 2;33(9):981–93. doi:10.1002/EMBJ.201488411/FIGURES/5 PubMed PMID: 24705786.

8. Fritsch C, Herrmann A, Nothnagel M, Szafranski K, Huse K, Schumann F, Schreiber S, Platzer M, Krawczak M, Hampe J, Brosch M. Genome-wide search for novel human uORFs and N-terminal protein extensions using ribosomal footprinting. Genome Res. 2012 Nov;22(11):2208–18. doi:10.1101/GR.139568.112 PubMed PMID: 22879431.

9. Xiao Z, Huang R, Xing X, Chen Y, Deng H, Yang X. De novo annotation and characterization of the translatome with ribosome profiling data. Nucleic Acids Res. 2018 Jun 1;46(10):E61. doi:10.1093/NAR/GKY179 PubMed PMID: 29538776.

10. Ji Z. RibORF: Identifying genome-wide translated open reading frames using ribosome profiling. Curr Protoc Mol Biol. 2018 Oct 1;124(1):e67. doi:10.1002/CPMB.67 PubMed PMID: 30178897.

11. Zhang P, He D, Xu Y, Hou J, Pan BF, Wang Y, Liu T, Davis CM, Ehli EA, Tan L, Zhou F, Hu J, Yu Y, Chen X, Nguyen TM, Rosen JM, Hawke DH, Ji Z, Chen Y. Genome-wide identification and differential analysis of translational initiation. Nature Communications 2017 8:1. 2017 Nov 23;8(1):1749-. doi:10.1038/s41467-017-01981-8 PubMed PMID: 29170441.

12. Guttman M, Russell P, Ingolia NT, Weissman JS, Lander ES. Ribosome profiling provides evidence that large noncoding RNAs do not encode proteins. Cell. 2013 Jul 3;154(1):240–51. doi:10.1016/j.cell.2013.06.009 PubMed PMID: 23810193.

13. Ingolia NT. Ribosome profiling: new views of translation, from single codons to genome scale. Nature Reviews Genetics 2014 15:3. 2014 Jan 28;15(3):205–13. doi:10.1038/nrg3645 PubMed PMID: 24468696.

14. Dunham I, Kundaje A, Aldred SF, Collins PJ, Davis CA, Doyle F, Epstein CB, Frietze S, Harrow J, Kaul R, Khatun J, Lajoie BR, Landt SG, Lee BK, Pauli F, Rosenbloom KR, Sabo P, Safi A, Sanyal A, Shoresh N, Simon JM, Song L, Trinklein ND, Altshuler RC, Birney E, Brown JB, Cheng C, Djebali S, Dong X, Ernst J, Furey TS, Gerstein M, Giardine B, Greven M, Hardison RC, Harris RS, Herrero J, Hoffman MM, Iyer S, Kellis M, Kheradpour P, Lassmann T, Li Q, Lin X, Marinov GK, Merkel A, Mortazavi A, Parker SCJ, Reddy TE, Rozowsky J, Schlesinger F, Thurman RE, Wang J, Ward LD, Whitfield TW, Wilder SP, Wu W, Xi HS, Yip KY, Zhuang J, Bernstein BE, Green ED, Gunter C, Snyder M, Pazin MJ, Lowdon RF, Dillon LAL, Adams LB, Kelly CJ, Zhang J, Wexler JR, Good PJ, Feingold EA, Crawford GE, Dekker J, Elnitski L, Farnham PJ, Giddings MC, Gingeras TR, Guigó R, Hubbard TJ, Kent WJ, Lieb JD, Margulies EH, Myers RM, Stamatoyannopoulos JA, Tenenbaum SA, Weng Z, White KP, Wold B, Yu Y, Wrobel J, Risk BA, Gunawardena HP, Kuiper HC, Maier CW, Xie L, Chen X, Mikkelsen TS, Gillespie S, Goren A, Ram O, Zhang X, Wang L, Issner R, Coyne MJ, Durham T, Ku M, Truong T, Eaton ML, Dobin A, Tanzer A, Lagarde J, Lin W, Xue C, Williams BA, Zaleski C, Röder M, Kokocinski F, Abdelhamid RF, Alioto T, Antoshechkin I, Baer MT, Batut P, Bell I, Bell K, Chakrabortty S, Chrast J, Curado J, Derrien T, Drenkow J, Dumais E, Dumais J, Duttagupta R, Fastuca M, Fejes-Toth K, Ferreira P, Foissac S, Fullwood MJ, Gao H, Gonzalez D, Gordon A, Howald C, Jha S, Johnson R, Kapranov P, King B, Kingswood C, Li G, Luo OJ, Park E, Preall JB, Presaud K, Ribeca P, Robyr D, Ruan X, Sammeth M, Sandhu KS, Schaeffer L, See LH, Shahab A, Skancke J, Suzuki AM, Takahashi H, Tilgner H, Trout D, Walters N, Wang H, Hayashizaki Y, Reymond A, Antonarakis SE, Hannon GJ, Ruan Y, Carninci P, Sloan CA, Learned K, Malladi VS, Wong MC, Barber GP, Cline MS, Dreszer TR, Heitner SG, Karolchik D, Kirkup VM, Meyer LR, Long JC, Maddren M, Raney BJ, Grasfeder LL, Giresi PG, Battenhouse A, Sheffield NC, Showers KA, London D, Bhinge AA, Shestak C, Schaner MR, Kim SK, Zhang ZZ, Mieczkowski PA, Mieczkowska JO, Liu Z, McDaniell RM, Ni Y, Rashid NU, Kim MJ, Adar S, Zhang Z, Wang T, Winter D, Keefe D, Iyer VR, Zheng M, Wang P, Gertz J, Vielmetter J, Partridge EC, Varley KE, Gasper C, Bansal A, Pepke S, Jain P, Amrhein H, Bowling KM, Anaya M, Cross MK, Muratet MA, Newberry KM, McCue K, Nesmith AS, Fisher-Aylor KI, Pusey B, DeSalvo G, Parker SL, Balasubramanian S, Davis NS, Meadows SK, Eggleston T, Newberry JS, Levy SE, Absher DM, Wong WH, Blow MJ, Visel A, Pennachio LA, Petrykowska HM, Abyzov A, Aken B, Barrell D, Barson G, Berry A, Bignell A, Boychenko V, Bussotti G, Davidson C, Despacio-Reyes G, Diekhans M, Ezkurdia I, Frankish A, Gilbert J, Gonzalez JM, Griffiths E, Harte R, Hendrix DA, Hunt T, Jungreis I, Kay M, Khurana E, Leng J, Lin MF, Loveland J, Lu Z, Manthravadi D, Mariotti M, Mudge J, Mukherjee G, Notredame C, Pei B, Rodriguez JM, Saunders G, Sboner A, Searle S, Sisu C, Snow C, Steward C, Tapanari E, Tress ML, Van Baren MJ, Washietl S, Wilming L, Zadissa A, Zhang Z, Brent M, Haussler D, Valencia A, Addleman N, Alexander RP, Auerbach RK, Balasubramanian S, Bettinger K, Bhardwaj N, Boyle AP, Cao AR, Cayting P, Charos A, Cheng Y, Eastman C, Euskirchen G, Fleming JD, Grubert F, Habegger L, Hariharan M, Harmanci A, Iyengar S, Jin VX, Karczewski KJ, Kasowski M, Lacroute P, Lam H, Lamarre-Vincent N, Lian J, Lindahl-Allen M, Min R, Miotto B, Monahan H, Moqtaderi Z, Mu XJ, O’Geen H, Ouyang Z, Patacsil D, Raha D, Ramirez L, Reed B, Shi M, Slifer T, Witt H, Wu L, Xu X, Yan KK, Yang X, Struhl K, Weissman SM, Penalva LO, Karmakar S, Bhanvadia RR, Choudhury A, Domanus M, Ma L, Moran J, Victorsen A, Auer T, Centanin L, Eichenlaub M, Gruhl F, Heermann S, Hoeckendorf B, Inoue D, Kellner T, Kirchmaier S, Mueller C, Reinhardt R, Schertel L, Schneider S, Sinn R, Wittbrodt B, Wittbrodt J, Jain G, Balasundaram G, Bates DL, Byron R, Canfield TK, Diegel MJ, Dunn D, Ebersol AK, Frum T, Garg K, Gist E, Hansen RS, Boatman L, Haugen E, Humbert R, Johnson AK, Johnson EM, Kutyavin T V., Lee K, Lotakis D, Maurano MT, Neph SJ, Neri F V., Nguyen ED, Qu H, Reynolds AP, Roach V, Rynes E, Sanchez ME, Sandstrom RS, Shafer AO, Stergachis AB, Thomas S, Vernot B, Vierstra J, Vong S, Wang H, Weaver MA, Yan Y, Zhang M, Akey JM, Bender M, Dorschner MO, Groudine M, MacCoss MJ, Navas P, Stamatoyannopoulos G, Beal K, Brazma A, Flicek P, Johnson N, Lukk M, Luscombe NM, Sobral D, Vaquerizas JM, Batzoglou S, Sidow A, Hussami N, Kyriazopoulou-Panagiotopoulou S, Libbrecht MW, Schaub MA, Miller W, Bickel PJ, Banfai B, Boley NP, Huang H, Li JJ, Noble WS, Bilmes JA, Buske OJ, Sahu AD, Kharchenko P V., Park PJ, Baker D, Taylor J, Lochovsky L. An Integrated Encyclopedia of DNA Elements in the Human Genome. Nature. 2012 Sep 6;489(7414):57. doi:10.1038/NATURE11247 PubMed PMID: 22955616.

15. Landt SG, Marinov GK, Kundaje A. modENCODE consortia ChIP-seq guidelines and practices of the ENCODE and References License Commons Creative [Internet]. 2012. doi:10.1101/gr.136184.111

16. Pei B, Sisu C, Frankish A, Howald C, Habegger L, Mu XJ, Harte R, Balasubramanian S, Tanzer A, Diekhans M, Reymond A, Hubbard TJ, Harrow J, Gerstein MB. The GENCODE pseudogene resource. Genome Biology 2012 13:9. 2012 Sep 5;13(9):R51-. doi:10.1186/GB-2012-13-9-R51 PubMed PMID: 22951037.

17. Sisu C, Muir P, Frankish A, Fiddes I, Diekhans M, Thybert D, Odom DT, Flicek P, Keane TM, Hubbard T, Harrow J, Gerstein M. Transcriptional activity and strain-specific history of mouse pseudogenes. Nature Communications 2020 11:1. 2020 Jul 29;11(1):3695-. doi:10.1038/s41467-020-17157-w PubMed PMID: 32728065.

18. Poliseno L, Salmena L, Zhang J, Carver B, Haveman WJ, Pandolfi PP. A coding-independent function of gene and pseudogene mRNAs regulates tumour biology. Nature 2010 465:7301. 2010 Jun 24;465(7301):1033–8. doi:10.1038/nature09144 PubMed PMID: 20577206.

19. Johnsson P, Ackley A, Vidarsdottir L, Lui WO, Corcoran M, Grandér D, Morris K V. A pseudogene long-noncoding-RNA network regulates PTEN transcription and translation in human cells. Nature Structural & Molecular Biology 2013 20:4. 2013 Feb 24;20(4):440–6. doi:10.1038/nsmb.2516 PubMed PMID: 23435381.

20. Kearse MG, Wilusz JE. Non-AUG translation: a new start for protein synthesis in eukaryotes. Genes Dev. 2017 Sep 1;31(17):1717. doi:10.1101/GAD.305250.117 PubMed PMID: 28982758.

21. Ravi Kumar YS, Bhavana AR, Aziz A, Roy R, Reddy AM, Mahadevappa P, Verma B. A comprehensive review of ribosomal RNA-derived fragments (rRFs): Emerging insights and future directions. Int J Biol Macromol. 2025 Nov;330:148265. doi:10.1016/j.ijbiomac.2025.148265

22. Mehra S, Phadnis S, Y.S. RK, Singh A, Verma B. Epitranscriptomic regulation of host–virus interactions. Biochimica et Biophysica Acta (BBA) - Molecular Basis of Disease. 2027 Jan;1873(1):168423. doi:10.1016/j.bbadis.2026.168423

23. Olexiouk V, Van Criekinge W, Menschaert G. An update on sORFs.org: a repository of small ORFs identified by ribosome profiling. Nucleic Acids Res. 2018 Jan 4;46(D1):D497–502. doi:10.1093/NAR/GKX1130 PubMed PMID: 29140531.

24. Vanderperre B, Lucier JF, Bissonnette C, Motard J, Tremblay G, Vanderperre S, Wisztorski M, Salzet M, Boisvert FM, Roucou X. Direct Detection of Alternative Open Reading Frames Translation Products in Human Significantly Expands the Proteome. PLoS One. 2013 Aug 12;8(8):e70698. doi:10.1371/JOURNAL.PONE.0070698 PubMed PMID: 23950983.

25. Madhry D, Malvankar S, Phadnis S, Srivastava RK, Bhattacharyya S, Verma B. Synergistic correlation between host angiogenin and dengue virus replication. RNA Biol. 2023 Jan;20(1):805–16. doi:10.1080/15476286.2023.2264003 PubMed PMID: 37796112.

26. Singh A, Roy R, Singh K, Sokhal P, Afroj S, Phadnis S, Y.S RK, Verma B. Dengue Virus Life Cycle and Host Protein Interactions: Focus on RNA Binding Proteins and Therapeutic Advances. Rev Med Virol. 2025 Mar 6;35(2). doi:10.1002/rmv.70025

27. Pyronnet S, Pradayrol L, Sonenberg N. A cell cycle-dependent internal ribosome entry site. Mol Cell. 2000 Apr 1;5(4):607–16. doi:10.1016/S1097-2765(00)80240-3 PubMed PMID: 10882097.

28. Cornelis S, Bruynooghe Y, Denecker G, Van Huffel S, Tinton S, Beyaert R. Identification and characterization of a novel cell cycle-regulated internal ribosome entry site. Mol Cell. 2000 Apr 1;5(4):597–605. doi:10.1016/S1097-2765(00)80239-7 PubMed PMID: 10882096.

29. Nelson BR, Makarewich CA, Anderson DM, Winders BR, Troupes CD, Wu F, Reese AL, McAnally JR, Chen X, Kavalali ET, Cannon SC, Houser SR, Bassel-Duby R, Olson EN. A peptide encoded by a transcript annotated as long noncoding RNA enhances SERCA activity in muscle. Science (1979). 2016 Jan 15;351(6270):271–5. doi:10.1126/science.aad4076

30. Huang JZ, Chen M, Chen D, Gao XC, Zhu S, Huang H, Hu M, Zhu H, Yan GR. A Peptide Encoded by a Putative lncRNA HOXB-AS3 Suppresses Colon Cancer Growth. Mol Cell. 2017 Oct 5;68(1):171–184.e6. doi:10.1016/j.molcel.2017.09.015 PubMed PMID: 28985503.

31. D V, N Aniruddha, Yadav Bhavana. RiboWin [Internet]. 2026. Available from: https://github.com/Vaibhav-81124/RiboWin

32. Calviello L, Mukherjee N, Wyler E, Zauber H, Hirsekorn A, Selbach M, Landthaler M, Obermayer B, Ohler U. Detecting actively translated open reading frames in ribosome profiling data. Nat Methods. 2016 Feb 1;13(2):165–70. doi:10.1038/nmeth.3688 PubMed PMID: 26657557.

33. Xiao Z, Zou Q, Liu Y, Yang X. Genome-wide assessment of differential translations with ribosome profiling data. Nature Communications 2016 7:1. 2016 Apr 4;7(1):11194-. doi:10.1038/ncomms11194 PubMed PMID: 27041671.

34. Li Q, Brown JB, Huang H, Bickel PJ. Measuring reproducibility of high-throughput experiments. Ann Appl Stat. 2011 Sep 1;5(3). doi:10.1214/11-AOAS466

35. Severin J, Lizio M, Harshbarger J, Kawaji H, Daub CO, Hayashizaki Y, Bertin N, Forrest ARR, Beckhouse A, Wells C, Vijayan D, Mason E, Wolvetang E, Hitchens K, Le Cao KA, Nielsen L, Fearnley L, Kenna T. Interactive visualization and analysis of large-scale sequencing datasets using ZENBU. Nat Biotechnol. 2014 Mar 1;32(3):217–9. doi:10.1038/NBT.2840 PubMed PMID: 24727769.

36. Casper J, Speir ML, Raney BJ, Perez G, Nassar LR, Lee CM, Hinrichs AS, Gonzalez JN, Fischer C, Diekhans M, Clawson H, Benet-Pages A, Barber GP, Vaske CJ, van Baren MJ, Wang K, Rodriguez YJP, Jenkins-Kiefer JA, Chalamala M, Haussler D, Kent WJ, Haeussler M. The UCSC Genome Browser database: 2026 update. Nucleic Acids Res. 2026 Jan 6;54(D1):D1331–5. doi:10.1093/nar/gkaf1250

