## Supplementary material for "RiboWin: A computational framework to identify non-canonical open reading frames": supp doc.

**Supplementary information**

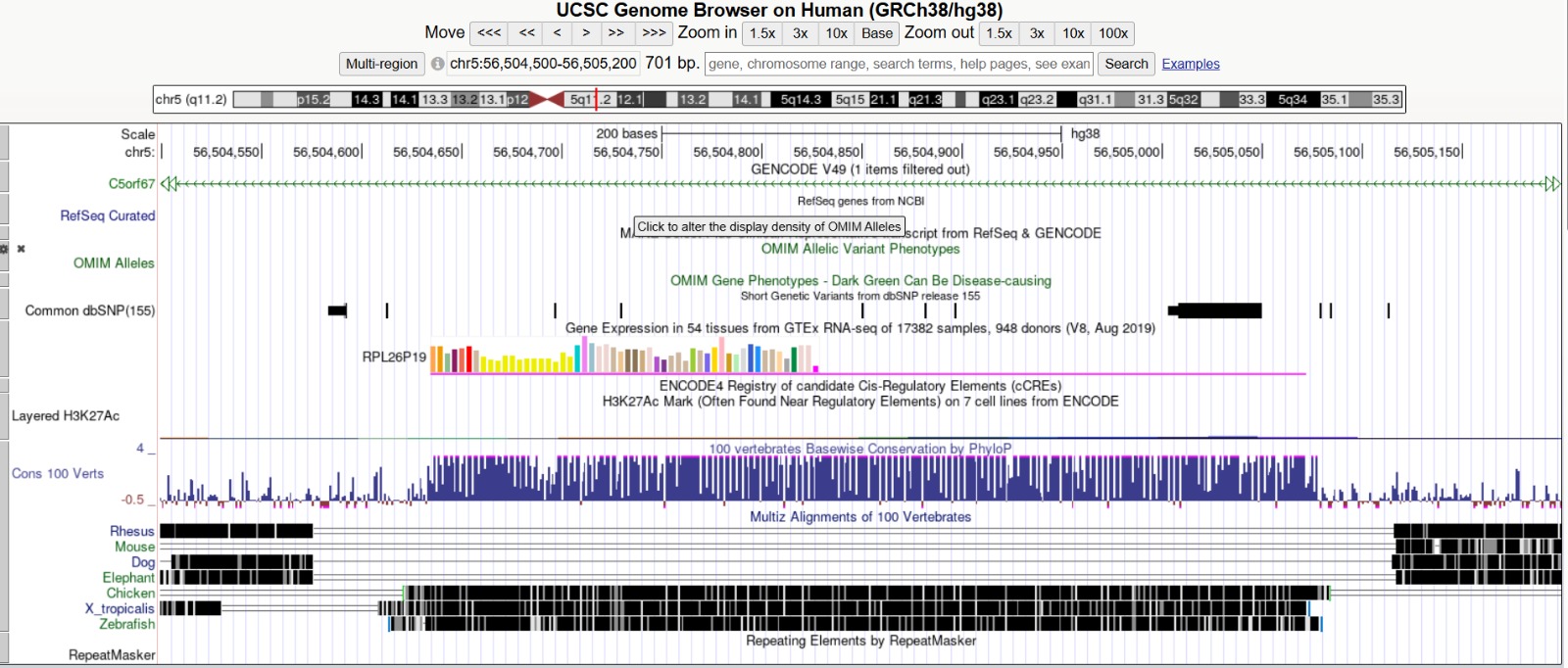

**S1**: UCSC Genome Browser view of the RPL26P19 locus (GRCh38). Gene annotation, GTEx expression, ENCODE regulatory elements (H3K27ac), evolutionary conservation, and repetitive regions can be observed.

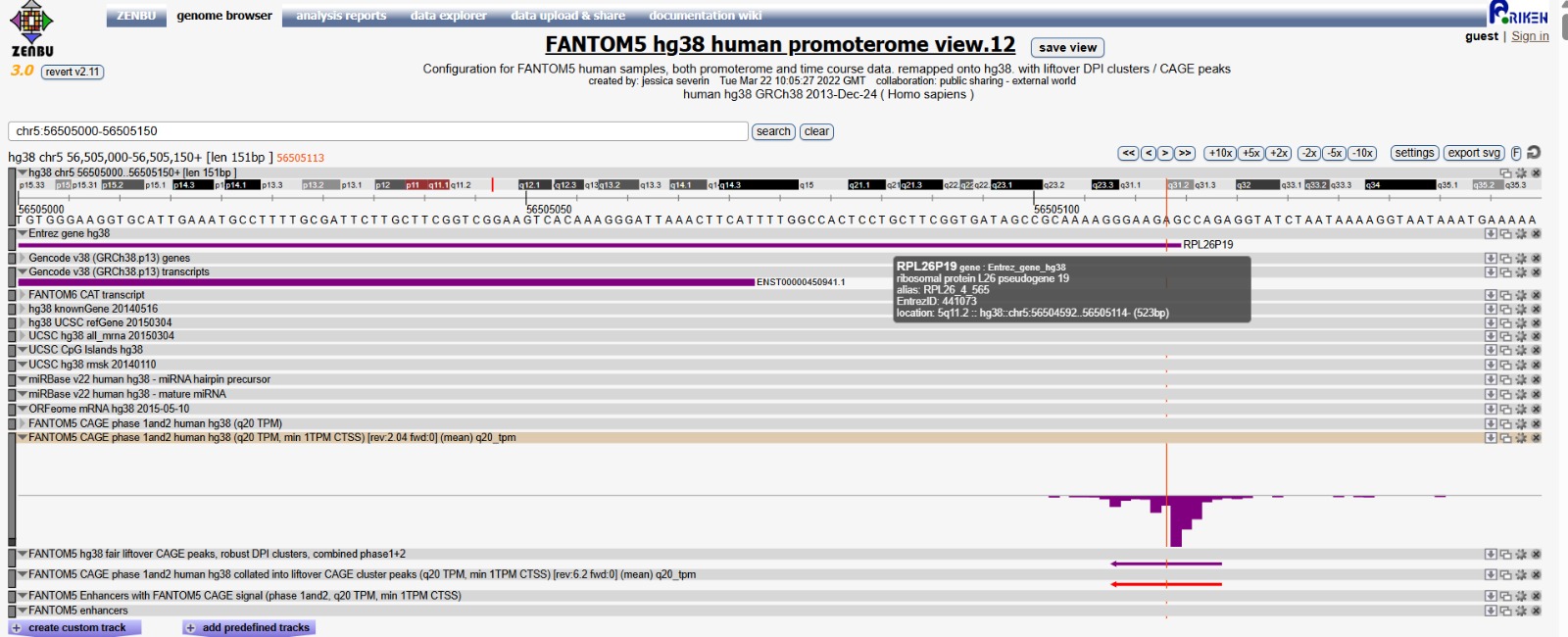

**S2**: FANTOM5 CAGE analysis of the RPL26P19 locus (GRCh38). FANTOM5 CAGE peaks identify transcription start site (TSS) activity near the 5′ end of RPL26P19, supporting promoter-associated transcriptional activity.

(S1: Supplementary information 1; S2: Supplementary information 2)
